# ARHGEF18 is a flow-responding exchange factor controlling endothelial tight junctions and vascular leakage

**DOI:** 10.1101/2022.09.10.507283

**Authors:** Surya Prakash Rao Batta, Marc Rio, Corentin Lebot, Céline Baron-Menguy, Reda Moutaoukil, Robin Le Ruz, Gervaise Loirand, Anne-Clémence Vion

## Abstract

The shear stress resulting from blood flow is a major regulator of endothelial cell (EC) biology and morphology. Rho protein-mediated cytoskeleton remodeling is an early and essential step of EC responses to flow. However, how Rho protein signaling is controlled by shear stress remains unclear. Here we demonstrate that phosphorylation, activity and expression of the Rho nucleotide exchange factor (RhoGEF) ARHGEF18 in ECs are modulated by the magnitude of shear stress. ARHGEF18 interacts with tight junctions, participates in ECs elongation and alignment and allows the maintenance of the endothelial barrier under physiological flow conditions. ARHGEF18 also promotes EC adhesion and migration by controlling focal adhesion formation. *In vivo*, ARHGEF18 is involved in flow response of ECs and in the control of vascular permeability in mice. Together, our results identified ARHGEF18 as the first flow-sensitive RhoGEF in ECs, whose activity is essential for the maintenance of intercellular junctions and the control of vascular permeability *in vivo*.

## Introduction

Mechano-sensitivity and mechano-transduction play crucial roles in shaping life. Cells, individually or collectively, need to respond appropriately to mechanical cues coming from their environment to adapt, survive, and to ensure healthy tissue development and homeostasis^1^. In the vascular system, endothelial cells (ECs) are exposed constantly to mechanical strains exerted by blood flow^2,3^. ECs are thus able to sense small variations in the direction, magnitude, and regularity of blood flow-induced shear stress (SS)^4,5^ and consequently modulate their functions to generate adapted responses. This leads, for instance, to arteriogenesis^6^, vasculature patterning^7,8^ and acute regulation of vessel tone,^9^ which are critical for both the development and maintenance of a healthy cardiovascular system. On the contrary, failure of ECs to adapt to changes in blood flow or chronic exposure to abnormal flow causes vascular diseases such as hypertension, atherosclerosis^3,10,11^ or intracranial aneurysm^12^.

Shear stress generated by laminar flow as observed in linear parts of the vascular tree is essential for the alignment of ECs in the direction of flow and the control of their functions^13,14^ thanks to several mechano-sensing mechanisms. Well-studied examples are ion channels^15^, primary cilia expressed at the apical surface of cells^16,17^, but also include mechano-sensing complexes at cell-cell interface such as the PECAM-1/VEGFRs/VE-cadherin complexe^10^ and at cell membrane-extracellular matrix attachment such as focal adhesions and integrins^18^. These sensors and complexes play an essential role in transforming flow-induced physiological forces into intracellular signals resulting in various cellular responses such as chemokine release, nitric oxide production, cell proliferation and junction reorganization. Rho protein-mediated actin cytoskeleton reorganization is one of the first steps of these processes^19–23^. Exposure of EC to physiological SS rapidly induced the translocation of Rho proteins from the cytosol to the plasma membrane associated with their activation^24^. Indeed, the overexpression of dominant-negative mutants of RhoA or Rac1 in ECs prevents their proper elongation and alignment in response to flow^25^. Both transient^25,26,13,18^ and sustained^24^ activation of Rho proteins upon SS are necessary for the establishment and maintenance of SS-induced EC alignment. Activation of Rho proteins consists in their transition from an inactive state (GDP-bound) to an active state (GTP-bound)^27^. By promoting the release of GDP followed by GTP loading, the guanine nucleotide exchange factors of Rho proteins (RhoGEFs) induce Rho protein activation^28^. In contrast to the ubiquity of Rho proteins, RhoGEF expression is tissue-specific and their activation is signal/time-specific^29^, placing them at the center of the control of the multiple effects of Rho proteins on cell functions. Some RhoGEFs such as FGD5^30^, have been described to be exclusively expressed in ECs, while others, although not specific to ECs, have been characterized for their role in mechano-sensitivity of ECs. The RhoGEFs Tiam1 and Vav2 are involved in the rapid activation of Rac1 in EC exposed to SS^31^. The RhoGEF Trio keeps Rac1 active at the downstream side of the EC and is essential for long-term flow-induced cell alignment^32^ as well as barrier integrity through the formation of a NOTCH1/VE-Cadh/LAR/Trio complex^33^. Lastly, ARHGEF12 has been identified as the RhoGEF responsible for RhoA activation in response to force detection by ICAM in EC promoting leukocyte transmigration^34^.

Here, using unbiased RNA screening in ECs, we identified ARHGEF18 as a new flow-sensitive RhoGEF. We demonstrated that ARHGEF18 is a RhoA-specific RhoGEF interacting with ZO1 and Claudin5 in ECs. We showed that ARHGEF18 participates in ECs alignment in response to flow by controlling p38 activity, tight junction and focal adhesion formation through its guanine nucleotide exchange activity. *In vivo*, endothelial-specific deletion of ARHGEF18 in mice led to retinal and brain hemorrhages consistent with its involvement in endothelial junction stability.

## Results

### Shear stress regulates ARHGEF18 expression and activity

In order to identify RhoGEF expression modulated by SS, we performed a 3’RNA profiling on ECs exposed to different SS levels: no flow (static, 0 dyn/cm^2^); pathological low SS (LSS, 36 dyn/cm^2^); physiological SS (PSS, 16 dyn/cm^2^) and pathological high SS (HSS, 36 dyn/cm^2^). The observed increased expression of the flow-dependent gene *KLF2* with increased level of SS validated our experimental conditions (Fig 1A, Fig suppl 1B). Among the genes with differential expression between static and SS, we identified several RhoGEFs whose expression was dependent on SS level (Fig 1A and suppl 1A). Among them, only *ARHGEF18* displayed an expression inversely correlated with the level of SS (suppl 1A). This inverse correlation between ARHGEF18 mRNA expression and SS levels was further confirmed by qRT-PCR analysis (Fig 1B). At the protein level, ARHGEF18 expression increased under pathological SS conditions (LSS and HSS) compared to physiological condition (PSS) (Fig 1C). This difference in the expression pattern between ARHGEF18 mRNA and protein suggests that additional post-transcriptional effect of SS could affect ARHGEF18 protein level. We next assessed whether the nucleotide exchange activity of ARHGEF18 on RhoA and Rac1 was also modulated by SS by performing pull-down assay using nucleotide-free RhoA^G17A^ and Rac1^G15A^ mutants to trap active RhoGEF^35^. Under static condition, we were able to detect ARHGEF18 among the proteins coprecipitated with RhoA^G17A^ indicating a basal nucleotide exchange activity of ARHGEF18 on RhoA in EC in the absence of flow (Fig 1D). Under flow conditions, the maximal association between ARHGEF18 and RhoA^G17A^ was detected in EC subjected to PSS, whereas it was lower under LSS and HSS conditions, indicating a high exchange activity of ARHGEF18 on RhoA under physiological flow condition that was reduced in pathological SS situations (Fig 1D). In contrast, we were unable to trap ARHGEF18 with Rac1^G15A^, regardless of SS conditions (Fig 1E), while the RacGEF Vav^36^ used as positive control was found among proteins coprecipitated with Rac^G15A^ (Fig suppl 1C). Taken together, these data show that ARHGEF18 is a RhoGEF specific for RhoA in ECs whose expression and activity are modulated by flow, with a maximal activity under physiological SS conditions.

**Figure 1:**
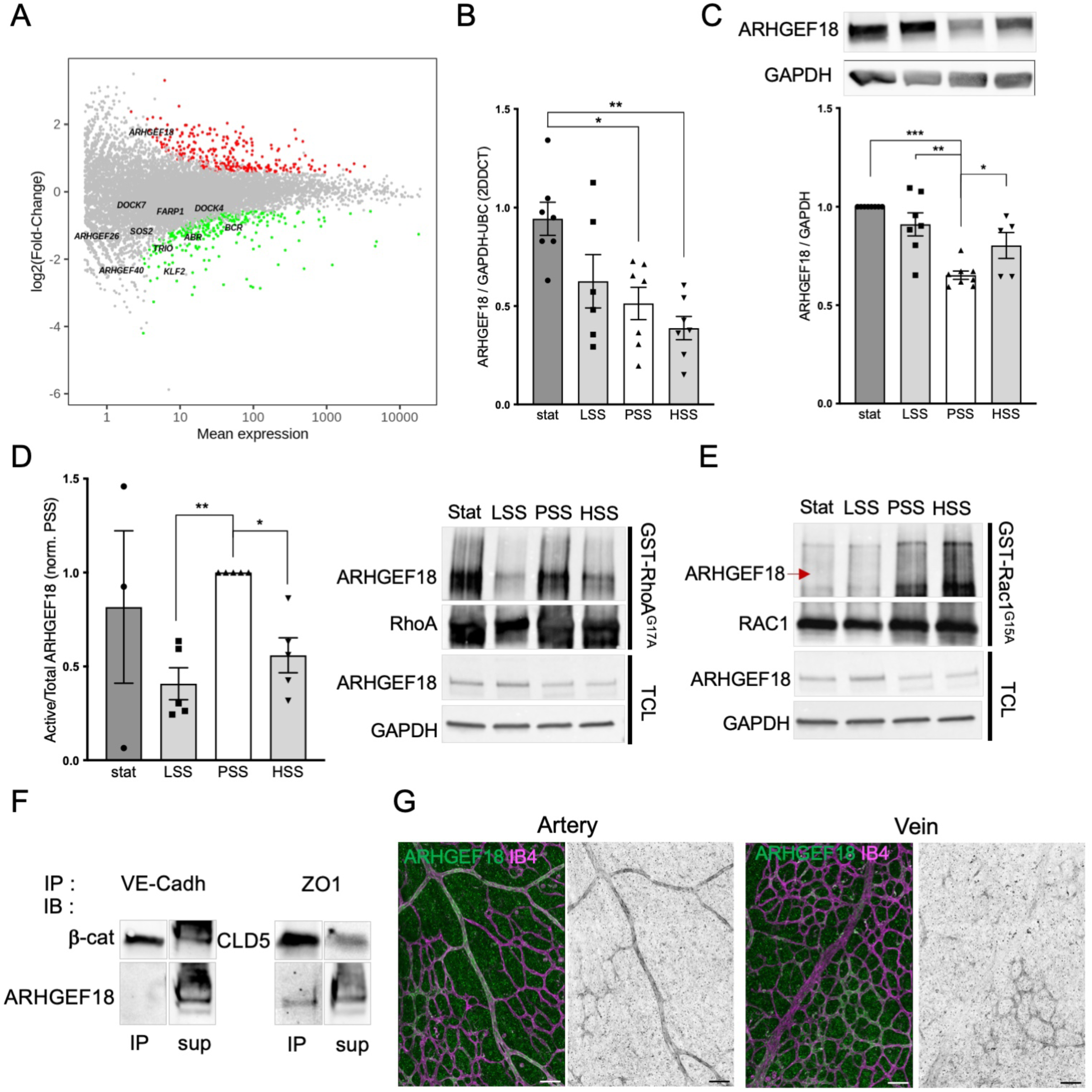
ARHGEF18 expression and activity is SS dependent. **A.** MA plot comparing gene expression in EC submitted for 24 hrs to static condition and HSS. Green dots: up-regulated genes in HSS condition; red dots: down-regulated genes in HSS condition. N=4. **B.** qPCR analysis of *ARHGEF18* mRNA expression level in EC exposed to static, LSS, PSS or HSS condition for 24 hrs. N>5. *p<0.05; **p<0.01; One-way ANOVA. **C.** Representative Western blot and quantification of ARHGEF18 protein expression level in EC exposed to static, LSS, PSS or HSS condition for 24 hrs. N>5. *p<0.05; **p<0.01; ***p<0.001; One-way ANOVA. **D.** Quantification and representative blot of ARHGEF18 activity determined from the amount of ARHGEF18 captured by GST-RhoA^G17A^ pull-down assay in EC exposed to static, LSS, PSS or HSS condition for 24 hrs. N=5. *p<0.05; **p<0.01; ***p<0.001; One-way ANOVA. **E.** Representative Western blot of ARHGEF18 captured by GST-Rac1^G15A^ pull-down assay in EC submitted to static, LSS, PSS or HSS condition for 24 hrs (representative of 3 independent experiment). **F.** Representative Western blot from Co-immunoprecipitation of ARHGEF18 with VE-cadherin (left) or ZO-1 (right). Co-immunoprecipitation efficiency was assessed by the presence of β-catenin and Claudin 5 in the immunoprecipitated fraction of VE-cadherin complex and ZO-1 complex respectively (upper lanes)(representative of 3 independent experiment). **G.** Immunofluorescent staining of ARHGEF18 in mouse retina from P6 animals. Green: Arhgef18, Magenta: IsolectinB4 (representative of 7 independent animals). Scale bar : 50μm.

### ARHGEF18 is expressed in arterial EC and interacts with tight junction proteins

ARHGEF18 has been shown to participate in tight junction formation in epithelial cells^37^. We thus assessed the possible interaction of endogenous ARHGEF18 with endothelial cell-cell junction proteins by co-immunoprecipitation experiments. ARHGEF18 was not detected in the protein fraction co-immunoprecipitated with VE-cadherin but was present in the immunoprecipitated fraction of the ZO-1 complex (Fig 1F). We confirmed the association of ARHGEF18 and ZO-1 in a same protein complex by the detection of ZO-1 in the proteins co-immunoprecipitated by an anti-flag antibody from the lysates of ECs expressing a flag-tagged ARHGEF18 (Fig suppl 1D). We also detected large amount of the major EC tight junction protein claudin 5 in this immunoprecipitate, while VE-cadherin was not observed (Fig suppl 1D). These results are consistent with the localization of ARHGEF18 within the tight junction protein complex. We next used immunofluorescence staining of mouse retinas to characterize the expression of ARHGEF18 in the vasculature. Using isolectin B4 staining to label ECs, we observed ARHGEF18 expression in the arteries of the retina at post-natal day 6, and in some capillary areas but not in the veins (Fig 1G). In 4-week-old mouse retina, this arterial expression was maintained but the capillary cluster were lost (Fig suppl 1E). Interestingly the staining was stronger in arteries close to the optic nerve (Fig suppl 1F) and fainted as the arteries caliber decreased and as the network ramified to become capillary (Fig suppl 1F). Altogether, these data showed that ARHGEF18 is expressed in arterial EC *in vivo* and localizes to the tight junction by interacting with ZO-1.

### ARHGEF18 participates in ECs adhesion and migration

We used siRNA and shRNA silencing strategies to turn off the expression of ARHGEF18 (Fig suppl 2A and 2B) and analyzed its role in ECs. We first evaluated the consequence of ARHGEF18 depletion on RHOA activity by pull-down assay. Under PSS condition, activity of RHOA was decreased by 30% in ARHGEF18-depleted EC compared to controls (Fig suppl 2C). Surprisingly, activity of RAC1 was also decreased by 30% in ARHGEF18-depleted EC compared to controls (Fig suppl 2D) although ARHGEF18 not being a GEF for RAC1^38^. We next assessed the functional role of ARHGEF18 in ECs in cell processes that depend on the activity of RHO proteins and the dynamic organization of the actin cytoskeleton such as adhesion and migration. Real-time impedance measurement after EC seeding showed that ARHGEF18 depletion reduced both the slope of impedance rise and the plateau, indicating that ARHGEF18 regulates early adhesion and spreading of ECs (Fig 2 A-F). To further characterize this effect at the single-cell level, we plated ECs onto L-shaped micro-patterned fibronectin to overcome variability in cell morphology by forcing cells to individually adhere and adopt a normalized geometry, allowing a better comparison and quantification of the number and size of focal adhesions. Silencing of ARHGEF18 by siRNA and shRNA led to a significant decrease in the number of long focal adhesions (> 5 µm) compared to control conditions suggesting a defect in focal adhesion maturation in ARHGEF18- depleted ECs (Fig 2G and H). To assess the role of ARHGEF18 in EC migration, we used an *in vitro* wound healing assay which revealed that the closure of the wound was slowed down by ARHGEF18 knockdown (Fig 2I, J, K). These results indicate that ARHGEF18 promotes adhesion, focal adhesion sites formation and migration of ECs cultured without flow.

**Figure 2:**
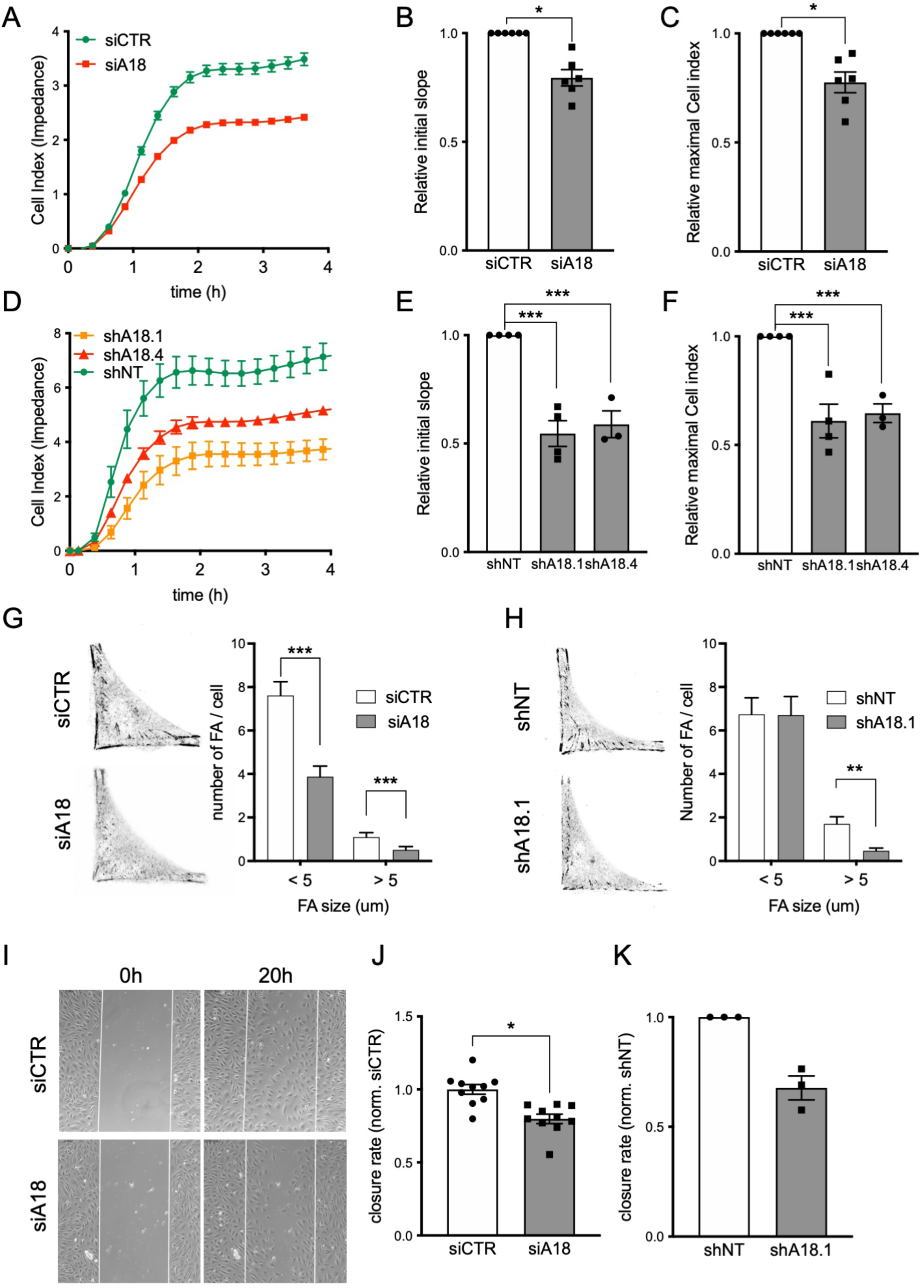
ARHGEF18 participates to endothelial cell adhesion and migration. **A.** Representative curves of impedance measurement during adhesion of HUVECs. Green: non-targeting siRNA (siCTR); red: siRNA targeting arhegf18 (siA18). **B.** Quantification of adhesion speed of HUVECs transfected with a control siRNA (siCTR) or with an siRNA targeting Arhgef18 (siA18). N=6. *p<0.05; Wilcoxon T-test. **C.** Quantification of maximal adhesion of HUVECs transfected with a control siRNA (siCTR) or with an siRNA targeting Arhgef18 (siA18). N=6. *p<0.05; Wilcoxon T-test. **D.** Representative curves of impedance measurement during adhesion of HUVECs. Green: non-targeting shRNA (shNT); red and orange: shRNAs targeting arhgef18 RNA (shA18.4 andshA18.1 respectively). **E.** Quantification of adhesion speed of HUVECs expressing a non-targeting shRNA (shNT) or a shRNA targeting Arhgef18 (shA18.1 or A18.4). N<3. ***p<0.001; One-way ANOVA. **F.** Quantification of maximal adhesion of HUVECs expressing a non-targeting shRNA (shNT) or a shRNA targeting Arhgef18 (shA18.1 or A18.4). N<3. ***p<0.001; One-way ANOVA. **G.** Representative images of HUVECs adhesion on L-shaped micropattern immune-stained for paxillin (siRNA transfected HUVECs) and Quantification of focal adhesion number and size. n=39 (cells) from 4 independent experiments. ***p<0.001; 2-way ANOVA. **H.** Representative images of HUVECs adhesion on L-shaped micropattern immune-stained for paxillin (shRNA expressing HUVECs) and Quantification of focal adhesion number and size. n=51 (cells) from 4 independent experiments. ***p<0.001; 2-way ANOVA. **I.** Representative images of wound assay experiment (phase contrast images) at starting point (T:0 hr) and 20 hrs post wound in siRNA transfected HUVECs. **J.** Quantification of closure speed of HUVECs transfected with a control siRNA (siCTR) or with an siRNA targeting Arhgef18 (siA18). N=10. *p<0.05; paired T-test. **K.** Quantification of closure speed of HUVECs expressing a control shRNA (shNT) or an Arhgef18 shRNA (shA18.1). N=3.

### ARHGEF18 participates in ECs response to flow

We next sought to assess whether the high ARHGEF18 activity in ECs under PSS condition contributes to the physiological response of ECs to flow. As expected, control ECs (shNT) aligned and elongated in the direction of flow. ECs harbored continuous VE-cadherin junction with junction-associated lamellipodia formation at the front and rear of the cells, continuous ZO-1 staining and cortical actin cables (Fig 3A, upper panel). By contrast, ARHGEF18-depleted ECs subjected to PSS failed to elongate and align in the flow direction as illustrated by the significant decrease in the flow response index (Fig 3A, lower panel, B and Fig suppl 3A, B). Additionally, ARHGEF18 silencing did not affect the proliferation rate of ECs under PSS, ruling out a possible effect of ECs density/packing on the cell shape change observed in ARHGEF18-depleted ECs submitted to PSS (Fig suppl 3C). Despite the presence of VE-cadherin at junction (Fig 3A), the shape of the adherent junctions seemed to be modified in ARHGEF18- deficient ECs compared to the control. The use of a classification method to evaluate junctional activity^39^ actually revealed that ARHGEF18-deficient ECs presented more active adherent junctions than control ECs (Fig 3C). Moreover, the number of gaps between ECs was significantly increased in ARHGEF18- deficient ECs compared to control (Fig 3A, D and Fig suppl 3A). ZO-1 localization at junction appeared disrupted and was significantly decreased by ARHGEF18 knock-down (Fig 3A, E; Fig suppl 3A). In the same line, cortical actin was significantly less abundant in ARHGEF18 deficient ECs compared to control (Fig 3A, F; Fig suppl 3A).

**Figure 3:**
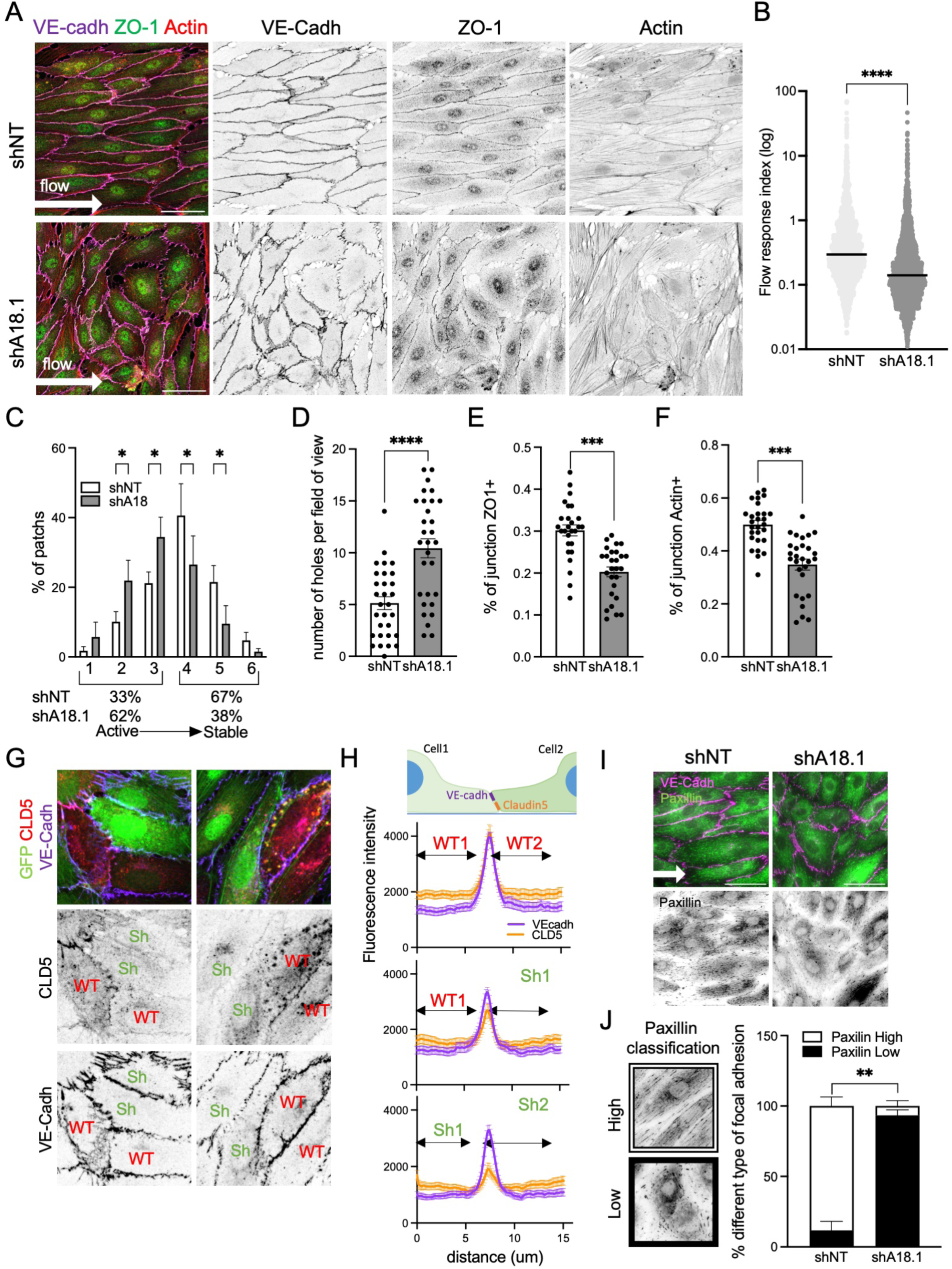
ARHGEF18 participates in ECs response to SS, tight junction formation and focal adhesion formation under physiological SS. **A.** Representative images of HUVECs expressing a non-targeting shRNA (shNT) or an Arhgef18 shRNA (shA18.1), exposed to physiological SS for 24 hrs. Flow direction is indicated by the arrow. Scale bar: 50μm **B.** Quantification of cell orientation and elongation with the flow in HUVECs expressing a non-targeting shRNA (shNT) or an Arhegef18 shRNA (shA18.1), exposed to physiological SS for 24 hrs. Each dot represents a cell. N=5 with in-between 200 and 600 cells analyzed by experiment. ***p<0.001; Mann-Whithney T-test. **C.** Quantification of junction status based on their morphology (N=4, 1000 patches analyzed per N). **D.** Quantification of number of holes in-between cells in HUVECs expressing a non-targeting shRNA (shNT) or an Arhgef18 shRNA (shA18.1), exposed to physiological SS for 24 hrs. N=3, 10 areas analyzed by experiment. ***p<0.001; Unpaired T-test. **E.** Quantification of ZO-1 localized at junction in HUVECs expressing a non-targeting shRNA (shNT) or an Arhgef18 shRNA (shA18.1), exposed to physiological SS for 24 hrs. N=3, 9 areas analyzed by experiment. ***p<0.001; Unpaired T-test. **F.** Quantification of Actin localized at junction in HUVECs expressing a non-targeting shRNA (shNT) or an Arhgef18 shRNA (shA18.1), exposed to physiological SS for 24 hrs. N=3, 9 areas analyzed by experiment. ***p<0.001; Unpaired T-test. **G.** Representative images of Claudin5 staining in a mix population of WT cells (GFP negative, indicated as “WT” on images) and cells expressing an Arhgef18 shRNA (shA18.1, GFP positive, green, indicated as “KO” on images), exposed to physiological SS. Green: GFP; red: Claudin5; Magenta: VE-cadherin. **H.** Quantification of the Claudin5 staining at junction. N=3, 12 couple of cells analyzed per experiment. **I.** Representative images of Paxillin staining in HUVECs expressing a non-targeting shRNA (shNT) or an Arhgef18 shRNA (shA18.1), exposed to physiological SS. Flow direction is indicated by the arrow. Green: paxillin; magenta: VE-cadherin. Scale bar: 50μm. **J.** Quantification of focal adhesion type within each cell based on the representative images for each category on the left (white box: paxillin high = numerous, elongated and aligned focal adhesions; black box: paxillin low = few, short and misaligned focal adhesions). N=4.

To further determine the role of ARHGEF18 on EC junction under PSS condition, we analyzed claudin 5 labeling at cell-cell junction, taking advantage of the mosaicism of ECs expressing ARHGEF18 shRNA, detectable by GFP labelling in EC monolayer. We observed that the maximal intensity of claudin 5 labeling at junction (colocalizing with VE-cadherin) between two ECs decreased when one of the two ECs expressed the ARHGEF18-shRNA. This junctional localization of claudin 5 was even more strongly reduced if the two ECs on either side of the junction expressed the ARHGEF18-shRNA (Fig 3G and 3H, Fig suppl 3D). This effect is reflected by an overall decrease in claudin 5 expression analyzed by western blot in ARHGEF18 deficient ECs subjected to PSS compared with control non-targeting shRNA cells (Fig suppl 3E). This suggests that the loss of ARHGEF18 expression leads to tight junction destabilization under PSS which could affect claudin 5 protein expression or stability. To analyze focal adhesions, also known to participate in ECs response to flow^21^, we performed immunofluorescent staining of paxillin on ECs exposed to PSS. Paxillin staining showed that focal adhesions, which appeared elongated and collectively oriented in control ECs, became more punctiform and disorganized in ARHGEF18-deficient ECs (Fig 3I, J and Fig suppl 3F, G). Surprisingly, we did not observe any change in overall FAK phosphorylation level assessed by western blot in ARHGEF18-deficient ECs compared to controls (Fig suppl 3H). Altogether, these data suggest that ARHGEF18 contributes to ECs response to physiological flow by promoting tight junction formation/stabilization, actin recruitment at cell-cell junction, and participating in focal adhesion sites organization.

### ARHGEF18 activity is essential for ECs response to flow through p38 signaling pathway

We wanted to explore the possible involvement of AKT, p38 MAPK and ERK1/2 signaling pathways, known to play a central role in flow-dependent focal adhesion formation and actin remodeling in ECs^40–44,45,46^, in ARHGEF18-mediated effects. Western-blot analysis showed that ERK1/2 and AKT phosphorylation levels were unaffected in ARHGEF18-deficient ECs compared to control ECs (Fig suppl 4A and 4B). In contrast, PSS-induced p38 phosphorylation was significantly reduced by ARHGEF18 silencing (Fig 4A; Fig suppl 4C) suggesting that ARHGEF18 participates in the activation of p38 by flow and that p38 could be involved in ARHGEF18-dependent effect on tight junction and focal adhesion in EC in physiological flow condition. Indeed, pharmacological inhibition of p38 by SB239063 (100 nM) reproduced the effects of ARHGEF18 depletion by inducing loss of cell alignment and elongation (Fig suppl 4D and E). P38 inhibition also induced delocalization of ZO-1 and actin fibers from junctions in ECs under PSS (Fig suppl 4D, F and G). This loss in junctional ZO-1 was not accompanied by an increase in the number of gaps between ECs, suggesting an additional role of ARHGEF18 on cell contact maintenance which would be independent of p38. Last, p38 inhibition in ECs under PSS led to the loss of focal adhesion organization compared to control as observed with ARHGEF18 depletion (Fig suppl 4H and I).

**Figure 4:**
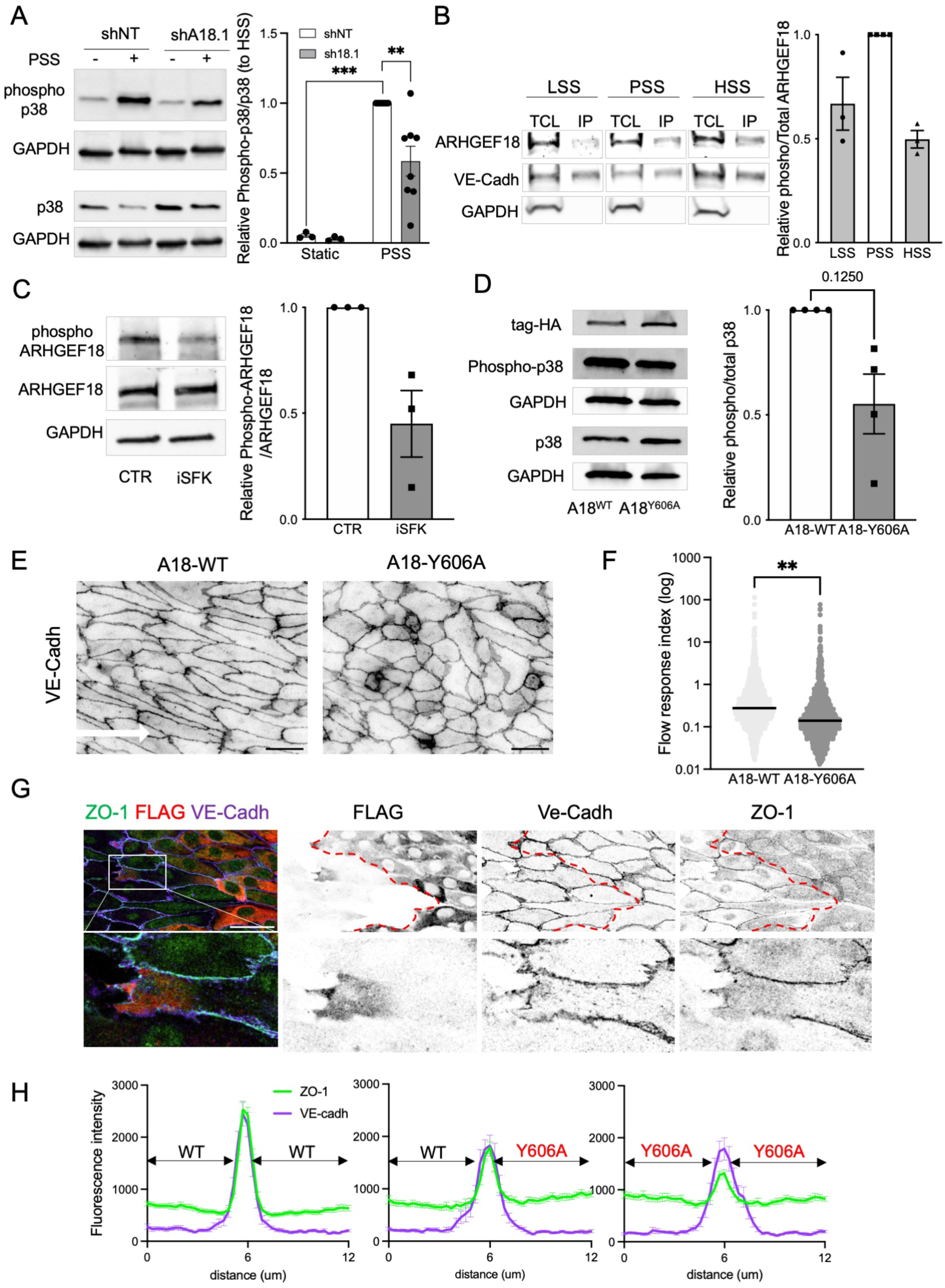
ARHGEF18 effects under flow are dependent of p38 and relies on its GEF activity. **A.** Representative images and quantification of p38 phosphorylation by Western blot in HUVECs expressing a non-targeting shRNA (shNT) or an Arhgef18 shRNA (shA18.1). Static: N=3; PSS: N=8. **p<0.01; ***p<0.001 2-way ANOVA**. B.** Representative images and quantification of ARHGEF18 Western blot from phosphor-tyrosin immunoprecipitation from HUVECs exposed to static, low SS (LSS), physiological SS (PSS) or high SS condition (HSS). N=3. **C.** Representative images and quantification of ARHGEF18 Western blot from phosphor-tyrosin immunoprecipitation from HUVECs exposed to PSS (3h) and treated with an inhibitor for SFKs (iSFK, SU6656) or vehicle (CTR), N=3. **D.** Representative images and quantification of p38 phosphorylation by Western blot in HUVECs expressing a WT form (A18^WT^) or a mutant form (A18^Y606A^) of ARHGEF18. **E.** Representative images of HUVECs exposed to physiological SS for 24 hrs, expressing a WT form (A18^WT^) or a mutant form (A18^Y606A^) of ARHGEF18 and stained for VE-cadherin. Flow direction is indicated by the arrow. Scale bar: 50μm. **F.** Quantification of cell orientation and elongation with the flow in HUVECs exposed to physiological SS for 24 hrs, expressing a WT form (A18^WT^) or a mutant form (A18^Y606A^) of ARHGEF18. Each dot represents a cell. between 1000 and 1350 cells analyzed. **p<0.01; Unpaired T-test. **G**. Representative images of ZO-1 staining in a mix population of WT cells (FLAG negative, non-red) and cells expressing the mutant form (A18^Y606A^) of ARHGEF18 (FLAG positive, red), exposed to physiological SS for 24 hrs. Green: ZO-1; red: FLAG; Magenta: VE-cadherin. Scale bar: 50μm. **H.** Quantification of the ZO-1 staining at junction. N=3, 25 couples of cells analyzed per experiment.

To further characterize the link between ARHGEF18, RHOA and p38, we sought to prevent the activation of ARHGEF18 by flow. By performing immuno-precipitation of phospho-tyrosinated proteins, we identified that ARHGEF18 was phosphorylated on tyrosine residues and this phosphorylation was increased in PSS conditions (Fig 4B). The phosphorylation of the tyrosine residue Y606 in the Dbl domain of ARHGEF18 has been shown to be necessary for its GEF activity^37^ and predicted to be under the control of a Src family kinase (SFK). Interestingly, the level of ARHGEF18 tyrosine phosphorylation under flow was strongly reduced by the SFK inhibitor SU6656 (Fig 4C), suggesting that flow-induced ARHGEF18 activation could depend on SFK-mediated phosphorylation of Y606. Based on this knowledge, we substituted this tyrosine with an alanine residue and expressed this phosphoresistant ARHGEF18 mutant (ARHGEF18^Y606A^) in ECs silenced for endogenous ARHGEF18. Pull-down assay with RHOA^G17A^ showed that, in contrast to WT-ARHGEF18, ARHGEF18^Y606A^ was not associated with RHOA under PSS condition, indicating a key role of Tyr606 phosphorylation in the GEF activity of ARHGEF18 under physiological flow conditions (Fig 4C). This loss of GEF activity in ARHGEF18^Y606A^- expressing ECs was associated with a decrease in p38 phosphorylation compared that of EC expressing wild-type (Fig 4D). Morphologically, expression of ARHGEF18^Y606A^ made ECs incapable of aligning in response to physiological flow, contrary to ECs expressing WT-ARHGEF18 (Fig 4E and F). To assess the consequence of ARHGEF18^Y606A^ expression on tight junction, we analyzed ZO-1 labeling in mosaic cells and again took advantage of the fact that not all ECs expressed the phosphoresisitant ARHGEF18 mutant, detectable by FLAG labelling. We observed that the intensity of ZO-1 labeling at the junction (colocalized with VE-cadherin) between two ECs decreased when one of the two neighboring ECs expressed ARHGEF18^Y606A^ compared to the intensity between two non-transfected ECs, and was further reduced when both ECs on either side of the junction expressed ARHGEF18^Y606A^ (Fig 4 G and H). Concurrently, the intracellular staining of ZO-1 was increased in EC expressing ARHGEF18^Y606A^ compared to control ECs (Fig G and H). Lastly, in ECs with local over-expression of the GEF-mutant, we observed a focal loss of ZO-1 staining at the membrane of the area where ARHGEF18^Y606A^ accumulated suggesting that ARHGEF18, through its GEF activity, could participate in the addressing of ZO-1 to the membrane (Fig 4G, lower panel). Overall, these data show that ARHGEF18 is activated by physiological flow through a mechanism involving Tyr606 phosphorylation, and that ARHGEF18 activation is required for p38 phosphorylation, tight junction organization and EC alignment under physiological flow condition.

### ARHGEF18 loss in vivo reduces vascular development and induces vascular leakage

To assess the function of ARHGEF18 *in vivo*, we generated tamoxifen inducible endothelial specific ARHGEF18 knockout mice (*Arhgef18^fl/fl^; CDH5-iCre^ERT2+/wt^* mice; referred to as *Arhgef18-iEC-KO*).The efficiency of tamoxifen treatment to induce ARHGEF18 deletion in ECs expressing CDH5-iCre and floxed Arhgef18 alleles *in vitro* was confirmed by western blot (Fig suppl 5A). In mice, analysis of Arhgef18 expression in lungs, used as an endothelium-rich tissue, showed a marked decrease in *Arhgef18*-iEC-KO, which also displayed a reduced expression of claudin 5 compared to lung from control littermates (*Arhgef18^fl/fl^* mice treated with tamoxifen, Fig suppl 5B). The retinal vasculature of P6 *Arhgef18*-iEC-KO mice showed a reduced radial expansion and a decreased vascular density compared with littermate controls (Fig 5A, B and C). This phenotype was transient since no more difference was observed between the retinal vasculature of 4-week-old *Arhgef18*-iEC-KO and littermate controls mice (data not shown). The reduced vascular density observed in the *Arhgef18*-iEC-KO retina (P6) was unlikely due to a reduced proliferation as the density of ECs present in the retinal vasculature was unchanged between *Arhgef18*-iEC-KO and littermate controls (Fig suppl 5D and E). We then assessed the consequence of Arhgef18 deletion on tight junction *in vivo* by the analysis ZO-1 localization. At P6, ZO-1 fluorescent labeling at junction was reduced in ECs of both retinal arteries and aorta of *Arhgef18*- iEC-KO mice compared to littermate control (Fig 5D, E and F). Interestingly, the loss of ZO-1 fluorescent labeling at junction was less visible in the veins and capillaries of the retina (Fig suppl F), in which ARHGEF18 is not detectable by immunofluorescence (Fig 1H), suggesting a link between ARHGEF18 expression and ZO-1 localization *in vivo*. We then used "en face" labelling of the aorta to assess ECs alignment, which depends on blood flow. Shape analysis of ECs showed a defect in cell elongation, quantified by the aspect ratio, in the aorta of *Arhgef18*-iEC-KO mice compared to littermate control, although the overall cell orientation remained in the direction of the flow (Fig 5G and H; Fig suppl 5G). Moreover, ECs of *Arhgef18*-iEC-KO mouse aorta displayed slightly but significantly more active junctions compared to control (Fig 5I). As tight junctions are associated with vascular barrier maintenance^47^, we assessed vascular leakage by looking at erythrocyte’s extravasation. *Arhgef18*-iEC- KO mouse retinas presented twice as many erythrocytes outside the vasculature than littermate control mouse retinas, both as diffused leakage with erythrocytes spread within the avascular space and as focal hemorrhages (Fig 6A, B and C). At P12, this leakage was still present but only as a diffuse leakage, without focal hemorrhages (Fig 6D and E) and was no longer observed in 4-week-old *Arhgef18*-iEC-KO mice (data not shown). This bleeding was not a side effect of the expression of the recombinase as retinas from *CDH5-iCre^ERT2+/wt^*mice without any flox allele did not display increased erythrocytes leakage nor decreased radial expansion compared to their littermate controls (Fig suppl 6A, B and C). The retinal vasculature being an extension of the brain vasculature with common developmental origins^48^, we also assessed potential vascular leakage in the brain of the *Arhgef18*-iEC-KO mice (P6). Interestingly, we observed small red spots at the surface of the brain in 50% of the *Arhgef18*-iEC-KO mice, but only in 10 % of the littermate controls (Fig 6F). To confirm this finding, we stained thick sections of the brain for erythrocytes and while we did not see any hemorrhage in control mice, we regularly found erythrocytes leakage in *Arhgef18*-iEC-KO mice (Fig 6G). Most of them were localized in the cortex in the immediate vicinity of penetrating arterioles. Interestingly we observed different features of bleeding at P6, with fresh bleeding visualize by accumulation of intact erythrocytes outside the vasculature (Fig 6G, *Arhgef18*-iEC-KO #1, red arrow) and resolving bleedings with dead and phagocyted erythrocytes (Fig 6G, *Arhgef18-*iEC-KO #2 and #3, red arrow). These data thus confirmed our in vitro results, showing the involvement of ARHGEF18 in flow response of ECs and in the control of vascular permeability *in vivo* in mice.

**Figure 5:**
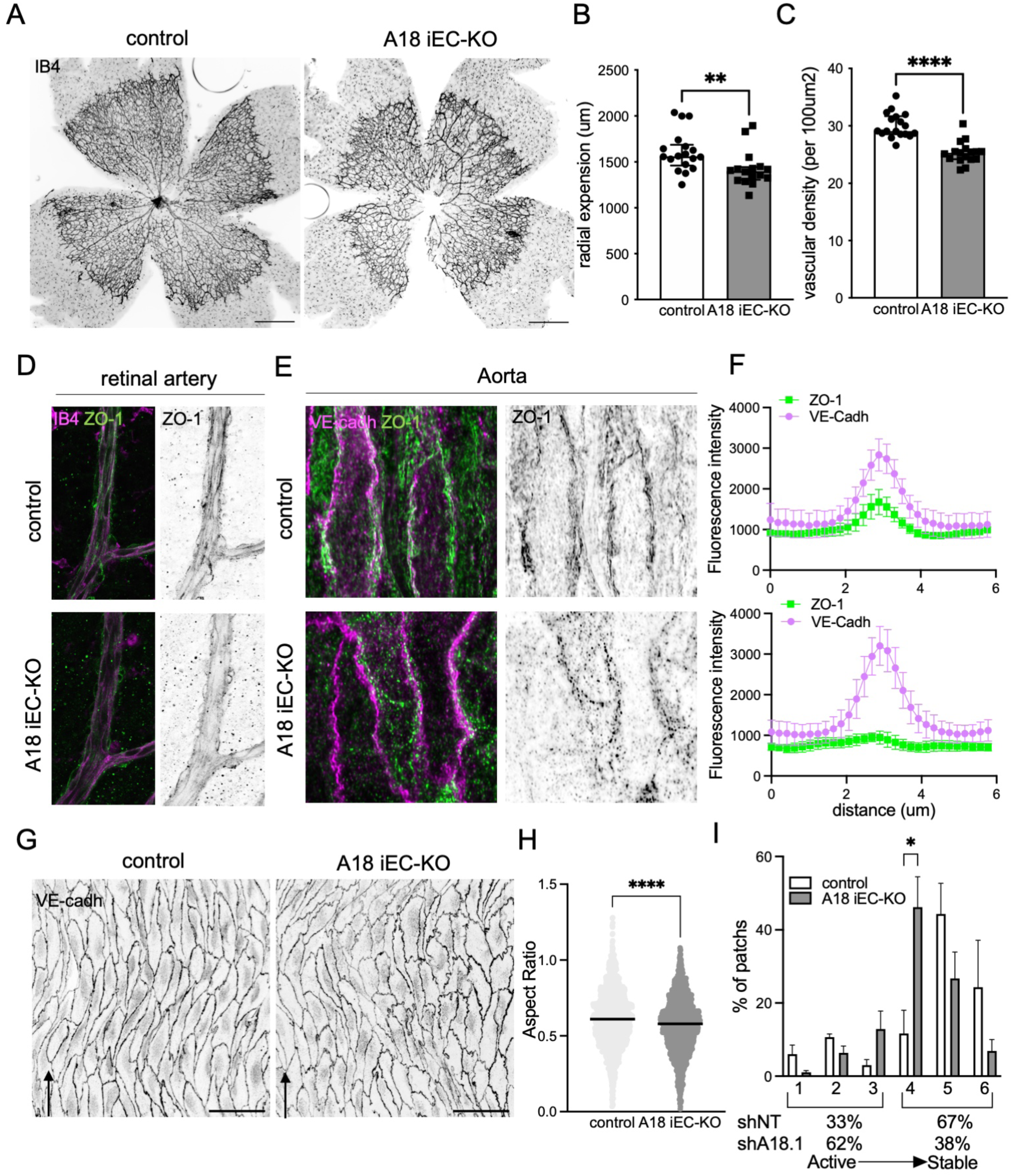
ARHGEF18 participates in vascular patterning and response to flow in vivo. **A.** Representative images of retinas from P6 pups wholemounts stained with Isolectin B4 (IB4, black). Scale bar: 500μm. **B.** Quantification of the radial expansion of the vascular plexus in *Arhgef18*-iEC-KO mice (A18 iEC-KO) and littermate controls (WT, n = 18; KO, n = 16; median ± IQ; Mann-Withney test). **C.** Quantification of the density of the vascular plexus in *Arhgef18*-iEC-KO mice and littermate controls (WT, n = 18; KO, n = 16; median ± IQ; unpaired-T test). **D.** Representative images of retinal artery stained for Isolectin (IB4, magenta) and ZO-1 (green) of P6 pups (observed in 3 littermate control and 4 *Arhgef18*-iEC-KO). **E.** Representative images of aortas mounted “en face” and stained for Isolectin (IB4, magenta) and ZO-1 (green) from P6 pups, imaged using Super-Resolution Structured Illumination Microscopy. **F.** Quantification of the ZO-1 staining at junction using plot profile analysis. N=3, 10 to 21 couples of cells analyzed per experiment. **G.** Representative images of aortas mounted “en face” and stained for VE-cadherin (gray) from 4 weeks-old mice. Scale bar: 50μm. **H.** Quantification of cell elongation (aspect ratio) with the flow in aortas. WT, N=6; *Arhgef18*-iEC-KO mice, N=7. Each dot represents a cell. with in-between 200 and 300 cells analyzed by experiment. ***p<0.001; unpaired T-test. **I.** Quantification of junction status based on their morphology (WT, N=3; *Arhgef18*-iEC-KO mice, N=6, 300 patches per N).

**Figure 6:**
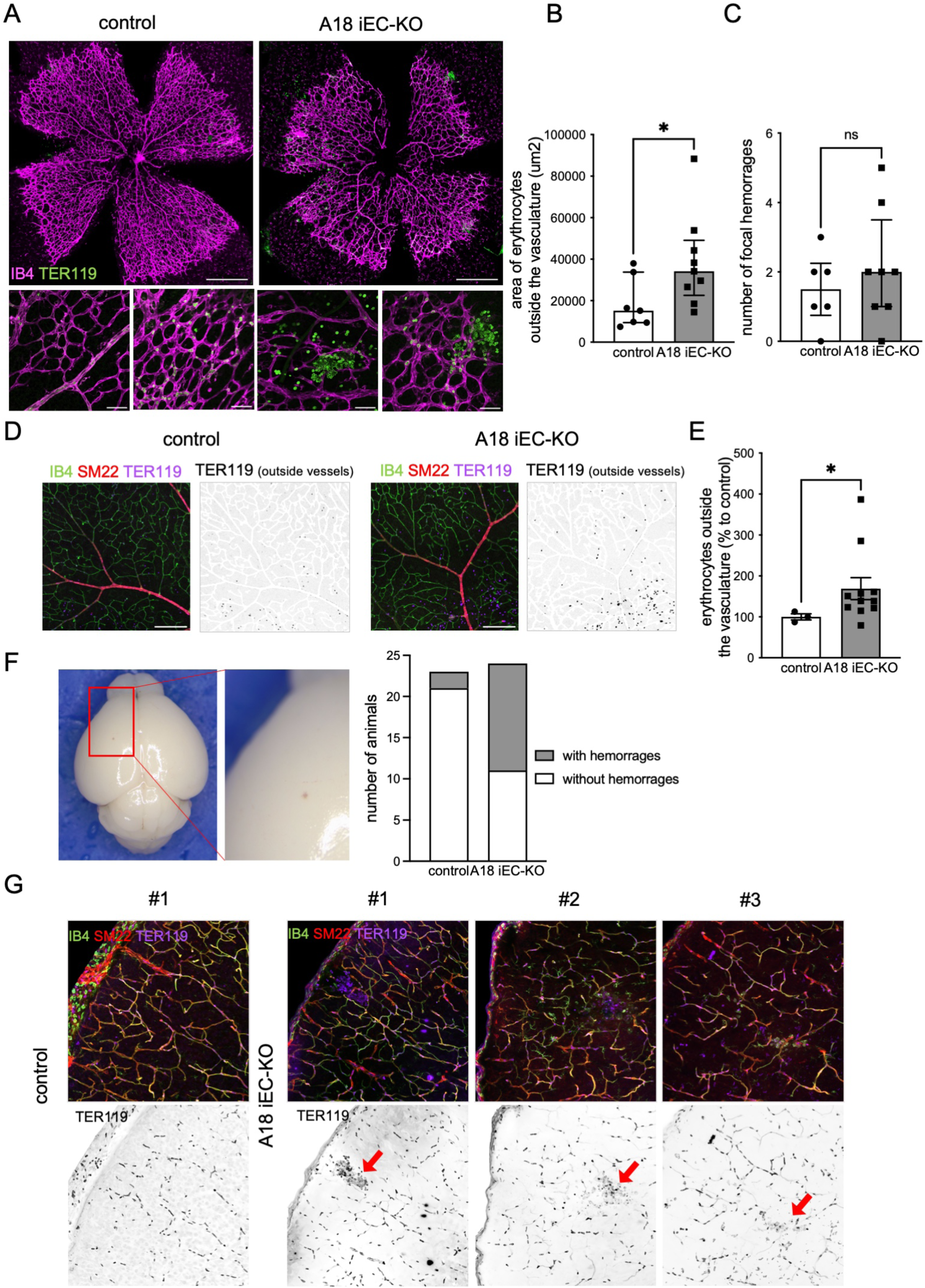
ARHGEF18 prevents retinas and brain hemorrhages. **A.** Representative retinal whole-mount images from P6 pups stained with isolectin (IB4, magenta) and TER119 (green). Scale bars: upper images, 500μm and lower images, 50μm. **B.** Quantification of erythrocytes outside the vasculature (control, N=7; *Arhgef18*-iEC-KO, N=9; median ± IQ; Mann-Withney T test). **C.** Quantification of leaking spot per retina (control, N=7; *Arhgef18*-iEC-KO, N=9; median ± IQ; Mann-Withney T test). **D.** Representative retinal whole-mount images from from P12 pups stained with isolectin (IB4, green), SM22 (red) and TER119 (violet). Scale bar: 200μm. **E.** Quantification of erythrocytes outside the vasculature (control, N=2; A18 iEC-KO, N=10; median ± IQ; Mann-Withney T test). **F.** Representative photography of brain from P6 animals and quantification of number of animals with micro-hemorrhages. **G.** Representative images of brain thick slices from P6 pups stained with isolectin (IB4, green), SM22 (red) and TER119 (violet). Red Arrow indicate leaking spots/ Representative of 3 WT and 4 KO.

## Discussion

Our present work identifies ARHGEF18 as a new SS-sensitive RhoGEF in ECs which participates in elongation and alignment of ECs and in maintenance of the endothelial barrier under physiological flow conditions. ARHGEF18, also known as p114-RhoGEF, is a RHOA-specific RhoGEF that is however able to trigger RAC1 activation in cells, probably through an indirect mechanism^38,49^. This is also true in ECs, wherein we showed that ARHGEF18 silencing was able to reduce RAC1 activity without a direct interaction. ARHGEF18 has been described mostly in epithelial cells where it controls tight junction maturation and apicobasal polarity establishment^37,50^ via RHOA and RAC1 activation^38,49,51^. ARHGEF18 activates RHOA at tight junctions, interacts directly with myosin IIA and regulates tight-junction assembly in a spatio-temporal manner, thus controlling long-range intercellular communication in epithelia and collective behavior of epithelial cells^37,52^. Here we demonstrate that ARHGEF18 is a mechano-sensitive protein in ECs: its expression, phosphorylation and activity are modulated by the magnitude of SS applied to ECs. ARHGEF18 activity is at its highest when ECs are exposed to physiological SS while its expression is at its lowest under these conditions. These observations are consistent with the existence of a negative regulatory loop between ARHGEF18 activity and expression. Indeed, activation of RHOA/RHO kinase signaling has been shown to generate a negative feedback loop on RHOA-GEF expression, including ARHGEF18^53^. This regulatory mechanism could therefore restrict ARHGEF18- induced RHOA activation by physiological flow, in agreement with the need of a moderate RHOA activity under physiological laminar flow to maintain EC alignment^26^.

It is well admitted that EC response to laminar flow is mediated by a rise in cytoskeletal tension that increases traction forces and intercellular tension, and promotes the assembly of adherens and tight junctions^54^ ^55^. As a major regulator of cytoskeletal reorganization, the dynamic regulation of RHOA is critical for SS-induced morphological reshaping of ECs^25,26^. Our results show that depletion of ARHGEF18 both *in vitro* and *in vivo* or expression of a catalytically inactive ARHGEF18 mutant impaired EC elongation and alignment with the flow in physiological conditions, thus indicating that ARHGEF18 is an essential component of the regulatory mechanisms that control RHOA activity and collective EC behavior of under flow.

The destabilization of tight junctions and junctional actin, and the associated increase in adherent junction activity in ARHGEF18-deficient ECs points toward a role of ARHGEF18 in sensing cell-cell tension arising from flow. This is consistent with the observed co-localization of ARHGEF18 with the tight junction proteins ZO-1 and claudin-5. Interestingly, ZO-1 has been recently shown to regulate tension on endothelial cell-cell contacts, in part via the recruitment of ARHGEF18 to a junctional protein complex^56^. Here we show that depletion of ARHGEF18 both *in vitro* and *in vivo* or expression of an inactive ARHGEF18 mutant compromised junctional recruitment of ZO-1, providing evidence that ARHGEF18 activity is mandatory to bring ZO-1 at tight junction and that ARHGEF18 and ZO-1 reciprocally need each other to localize at the junctional protein complex. Although ARHGEF18 does not co-localize with VE-cadherin, ARHGEF18 depletion also affects adherens junctions, without altering VE-cadherin localization. This last result confirms that ARHGEF18 is a key component of the molecular complex through which tight junction can indirectly regulate adherens junctions, as reported in EC under static conditions^56^. Interestingly, loss of ARHGEF18 not only disrupts tight junction under physiological SS but also impairs focal adhesion formation and organization, supporting the existence of a cross-talk between tight junction and focal adhesion. Since the generation and the transmission of force at cell-cell junctions and focal adhesions are important for the assembly and the distribution of adhesive complexes and junctions, this observation provides further evidence that the maintenance of junctional tension is an important function of ARHGEF18. This effect is likely to be the cause of the significant alteration of EC adhesion and migration induced by ARHGEF18 depletion *in vitro*, and participate to the reduced radial expansion of retinal arteries in *Arhgef18-*iEC-KO mice *in vivo*.

Our results obtained with the inactive ARHGEF18 mutant, together with the known role of RHOA in generating tension and in organizing EC layer in response to laminar flow strongly indicates that the effects of ARHGEF18 mostly depend on RHOA activation and that ARHGEF18 is responsible for laminar-flow-induced RHOA activation in ECs. Additionally, our data also identified p38 MAPK as a downstream effector of RHOA that mediates the effect of ARHGEF18 on tight junction and EC alignment under PSS conditions. Thus, while p38 MAPK activation has already been described for its role in flow-induced morphological changes in EC,^57,58^ our work identifies one of its upstream activator, ARHGEF18. We also partially decipher the mechanism upstream of ARHGEF18 activation by flow by showing that its activity depended on its phosphorylation on Y606, and that pharmacological inhibition of SFKs reduces its tyrosine phosphorylation level. According to the known differential efficiency of SU6656 to inhibit SKF members, the low concentration of SU6656 (50 nM) at which flow-induced phosphorylation of ARHGEF18 was inhibited is in favor of the involvement of the YES kinase^59^. This role of SFK, and more particularly YES, in ARHGEF18 activation is consistent with the impairment of PSS-induced EC alignment observed when SFK proteins are inhibited^58^ and the recent description of YES as a flow sensitive protein^60^.

In addition to RHOA/p38 signaling, other pathways likely contribute to the effect of ARHGEF18 since the intercellular gaps induced by ARHGEF18 depletion were not reproduced by expression of the inactive ARHGEF18 mutant or by p38 inhibition in EC under flow. This result supports the concept that RhoGEFs could regulate cell functions through scaffolding properties independent of their guanine nucleotide exchange activity on RHO proteins^61,62^.

In mice, complete deletion of ARHGEF18 resulted in embryonic lethality^63^. *Arhgef18* null embryos showed reduced vasculature, increased hemorrhage and edema, which is in line with our observation. Our mouse model, harboring EC specific deletion of ARHGEF18 after birth, presents bleedings in the retina and the brain associated with a loss of ZO-1 at junction. We did not observe spontaneous micro- hemorrhages outside de CNS. In the aortas, we also observed a mis-localization of ZO-1 which was not associated with vascular leakage. This vascular phenotype of *Arhgef18*-iEC-KO confirms *in vivo* the role of ARHGEF18 in the maintenance of tight junctions and EC alignment under physiological laminar flow and suggests a potential specificity of ARHEGF18 for the cerebral vasculature or a higher sensitivity to ARHGEF18 deletion in this area.

In humans, a variant of the *ARHGEF18* gene has been shown to be associated with systemic capillary leak syndrome^64^, a rare disorder characterized by repeated, transient, unprovoked episodes of hypotensive shock and peripheral edema due to a transient endothelial hyperpermeability, consistent with the role of ARHGEF18 we identified in this study. Mutations in *ARHGEF18* have been also reported in inherited retinal dystrophy, characterized by retina degeneration affecting photoreceptors, but patients carrying ARHGEF18 mutations also displayed attenuated retinal vessel density which could be in line with a primary effect of those mutations on vascular ECs^65^.

In summary, ARHGEF18 is a novel mechanosensitive RhoGEF activated by physiological SS that plays a central role in the regulation of tight junction, thereby driving the alignment of EC subjected to laminar flow and controlling vascular permeability.

## Material and Methods

### Mice

Mice were maintained at the “unité thérapeutique experimental” of Nantes university under standard husbandry conditions. The following mouse strains were used: C57/BL6J, *Arhgef18^lox/lox^*(generated from mice carrying the conditional knockout first *Arhgef18*^tm1a(KOMP)Mbp^ from the MRC), *CDH5(PAC)-CreERT2* (Kindly provided by Pr R.Adams). To generate endothelium specific *Arhgef18*-knock-out (KO) mice, we mated *Arhgef18^lox/lox^* to *CDH5(PAC)-CreERT2* transgenic mice to generate mice containing both the *CDH5(PAC)-CreERT2* allele and *Arhgef18* floxed alleles (*CDH5(PAC)-CreERT2*; *Arhgef18^lox/lox^* mice: *Arhgef18*-iEC-floxed mice). Cre-mediated deletion in EC (*Arhgef18*-iEC-KO mice) was induced by tamoxifen (T5648, Sigma) treatment (intraperitoneal injection of 20 µg/g animal at postnatal days 1, 2 and 3). Littermates tamoxifen-treated *Arhgef18^lox/lox^* mice were used as controls. Retinas, brain, lungs and aortas were collected at postnatal day 6 (P6), 12 (P12) or at 4 week-old (4WO) onwards. Animal procedures were performed in accordance with the animal license APAFIS #33434- 2021101214003205. The investigators were blinded to allocation during experiments.

### Cell culture and microfluidic chamber experiments

Human umbilical vein endothelial cells (HUVECs) (passage 2 to 6; PromoCell), chosen for their accessibility and well characterized response to flow, were routinely cultured in EBM media supplemented with the provided growth factors kit (Promocell). For flow experiments dedicated to pull-down, co-immunoprecipitation, or WB lysate collection, cells were cultured on 0.2% gelatin-coated slides (Menzel Glazer) and submitted to unidirectional laminar SS using peristaltic pumps (Gilson) connected to a glass reservoir (ELLIPSE) and the chamber containing the slide. For flow experiments dedicated immunofluorescence staining, cells were cultured on 0.2% gelatin-coated 0.4 ibidi slides (IBIDI) and unidirectional laminar SS was applied using the pumping system and control unit form IBIDI. Local SS was calculated using Poiseuille’s law and averaged to 3.6 dyn/cm^2^ (pathological low SS: LSS) 16 dyn/cm^2^ (physiological SS: PSS) or 36 dyn/cm^2^ (pathological high SS: HSS). For inhibition studies, confluent HUVEC were treated with either the p38 specific inhibitor (100 nM, SB239063; TOCRIS) for 1hr prior to physiological SS, or the SFK inhibitor SU6656 (40nM, TOCRIS) for 3h prior to physiological SS. Treatment was maintained during the flow stimulation.

### Mice EC isolation and tamoxifen treatment in vitro

Brain endothelial cells, isolated from *arhgef18^f/fl^*(Control) or *arhgef18^f/fl^; Cdh5-CreERT2* micecortical gray matter, were obtain through collagenase dispase digestion and BSA and Percoll density gradient centrifugation. Purified vessels were seeded onto collagen IV/fibronectin-coated dishes at high density (3 brain for 4cm^2^). Cells were then grown in EGM2-MV (promocell) with 5 μg/ml puromycin during the first 5 days ^66^. Brain ECs were then treated with hydroxy-tamoxifen (1ug/mL) for 3 days to induce recombination.

### 3’Sequencing RNA Profiling

Experimental protocol: The protocol was performed according to the 3′-digital gene expression (3′-DGE) approach developed by the Broad institute as described in https://doi.org/10.21203/rs.3.pex-1336/v1. The libraries were prepared from 10 ng of total RNA in 4µl. The mRNA poly(A) tails were tagged with universal adapters, well-specific barcodes and unique molecular identifiers (UMIs) during template-switching reverse transcription. Barcoded cDNAs from multiple samples were then pooled, amplified and tagmented using a transposon-fragmentation approach which enriches for 3’ends of cDNA : 200ng of full-length cDNAs were used as input to the Nextera™ DNA Flex Library Prep kit (ref #20018704, Illumina) and Nextera™ DNA CD Indexes (24 Indexes, 24 Samples) (ref #20018707, Illumina) according to the manufacturer’s protocol (Nextera DNA Flex Library Document, ref #1000000025416 v04, Illumina). Size library was controlled on 2200 Tape Station Sytem (Agilent Technologies). A library of 350-800 bp length was run on a NovaSeq 6000 using NovaSeq 6000 SP Reagent Kit 100 cycles (ref #20027464, Illumina) with 17*-8-105* cycles reads.

Bioinformatics protocol: Raw fastq pairs match the following criteria: the 16 bases of the first read correspond to 6 bases for a designed sample-specific barcode and 10 bases for a unique molecular identifier (UMI). The second read (104 bases) corresponded to the captured poly(A) RNAs sequence. Bioinformatics steps were performed using a snakemake pipeline (https://bio.tools/3SRP). Samples demultiplexing was performed with a python script. Raw paired-end fastq files were transformed into a single-end fastq file for each sample. Alignment on refseq reference transcriptome, available from the UCSC download site, was performed using bwa. Aligned reads were parsed and UMIs were counted for each gene in each sample to create an expression matrix containing the absolute abundance of mRNAs in all samples. Reads aligned on multiple genes or containing more than 3 mismatches with the reference were discarded. The expression matrix was normalized and differentially expressed genes (DEG) were searched using the R package deseq2 (doi: 10.1186/s13059-014-0550-8). If DEG were found, functional annotation was performed using the R package ClusterProfiler (doi: 10.1016/j.xinn.2021.100141 / doi: 10.1089/omi.2011.0118).

### Generation of Lentiviral vectors

For RNA interference, the sequence for non-target shRNA was previously described (Supplemental Table 1). For Arhgef18, specific sense and anti-sense DNA sequences were identified using free online services (http://sidirect2.rnai.jp/; http://web.stanford.edu/group/markkaylab/cgi-bin/)^67^. The sequences that met the optimal criteria were selected and synthesized as oligos in the form of miR-E backbone. The oligos were then amplified, cloned into LT3GEPIR and sequenced as previously described. The resultant lentiviral vector drives the expression of shRNA along with EGFP under TRE3G promoter, a doxycycline inducible promoter. For Arhgef18 overexpression constructs, Arhgef18 was amplified from HUVEC cDNA by overlap-extension PCR using the oligos and primers in Supplemental Table 2. The resultant amplicon was digested and clone into LT3(AsiSI)N1-GPIR (modified LT3GEPIR) vector in between AsiSI and EcoRI restriction sites. The resultant vector LT3N1-GPIR-Arhgef18 encodes Arhgef18-EGFP fusion protein under TRE3G promoter (WT-ARHGEF18). GEF mutant version of Arhgef18 was generated by site directed mutagenesis of tyrosine to alanine at amino acid position 606 of the canonical sequence (Q6ZSZ5-4 isoform) using primers A18-Y606A-SDM-Fr (5’-CATAACCAAAGCCCCAGTGCTGGTG-3’), A18-Y606A-SDM-Rev (5’-CGTTGTGTAACCAGGAGAATG-3’) and Q5 site directed mutagenesis kit (NEB E0554). This mutation corresponds to the Y260A mutation described by Terry and colleague^37^ and Acharya and colleague^68^ based on the Q6ZSZ5-1 isoform sequence. This LT3N1-GPIR-Arhgef18 vectors were further modified to substitute EGFP with 3XHA-FLAG tag, shArhgef18.6 to target endogenous Arhgef18 and P2A sequence to replace IRES using standard molecular cloning techniques. The resultant vector, Arhgef18- HA/FLAG_shArhgef18.6, is designated as ARHGEF18^Y606A^ in the main text.

### Lentivirus production, transduction and induction of gene of interest vectors

Lentivirus vector carrying either the shRNA or Arhgef18 ORF was transfected into HEK293T cells along with packaging vectors psPAX2 and pVSVG2 (provided by Prof. Dr.Utz Fischer’s lab) using polyethylenimine (764965; Sigma-Aldrich) transfection agent. After overnight incubation, the medium was replaced with fresh medium (DMEM+10%serum+PenStrep). Forty-eight hrs and 72 hrs after transfection, viral supernatant was collected, filtered through 0.22 µm filter and used to infect HUVECs (passage 2) in the presence of polybrene (8 µg/ml, H9268; Sigma-Aldrich). The transduced HUVECs were selected with puromycin (8ug/ml, P8833; Sigma-Aldrich) for 48 hrs and maintained in the presence of puromycin (1µg/ml). The induction of either shRNA or full length ARHGEF18 was performed at least 2 days before the experiment by supplementing the medium with doxycycline (1µg/ml, D9891; Sigma-Aldrich) and maintained throughout the experimental procedure.

### Pulldown Assay

Pulldown of active small GTPases (RHOA/RAC) or active ARHGEF18 was done as previously described^1,2^. Briefly, HUVECs (WT, shRNA or A18-Y606A) subjected to various SS conditions for indicated time, were lysed in either RBD lysis buffer (50 mM Tris-HCl (pH 7.6), 500 mM NaCl, 1% (vol/vol) Triton X-100, 0.1% (wt/vol) SDS, 0.5% (wt/vol) deoxycholate and 10 mM MgCl2) or GEF lysis buffer (20 mM HEPES pH 7.5, 150 mM NaCl, 5 mM MgCl_2_, 1% (vol/vol) Triton X-100, 1 mM DTT) supplemented with protease and phosphatase inhibitor cocktail. Cell debris was removed by centrifugation and equal volumes of cell lysate was incubated with 25-50 ug of either GST-RBD/PBD for active small-GTPases; or GST tagged RHOA^G17A^/RAC1^G15A^ for active GEF pulldown for 45 min. After incubation, beads were washed three time in respective wash buffers. Captured proteins were detected using Western Blot.

### Immunoprecipitation and Co-immunoprecipitation assay

HUVECs subjected to various SS conditions for indicated time, were lysed in IP lysis buffer (1% Nonidet P-40, 25 mM Tris-HCl pH 7.4, 100 mM NaCl, 10 mM MgCl2, 10% glycerol) supplemented with protease and phosphatase inhibitor cocktail. Lysis buffer was homogenized passing through a 32-gauge syringe needle and cellular debris were removed by centrifugation (5 min, 10000 rpm). A small amount of the lysate was collected in order to assess the presence of the protein in the total cell lysate. Beads were first equilibrated in IP buffer and then coated with 0.5 to 1 µg of the following antibodies: phospho-Tyr (CST 9411), ZO-1 (Thermofisher), VE-cadherin (R&D systems) or FLAG (Merck, F1804). Cell lysats were then incubated with the coated beads, supernatants and immunoprecipitats (washed 3 times) were stored to proceed for Western blot.

### Micropatterning assay

Pattern shape was design with FIJI software to have a L shaped pattern 50x50 µm long and 6 µm large. PDMS stencils were placed in IBIDI wells and cleaned in a Plasma cleaner for 5 min. A hydrophobic treatment was done using PLL PEG (Poly-lysine Polyethylene glycol 100µg/ml) for 30min. Patterns were then prepared using the Primo device (Alveole). Briefly, the wells with PLPP (photoinitiator, Alveole) were placed under a microscope with the PRIMO device. Focus was adjusted to UV reflection and the patterning sequence was launched with 1200mJ/mm2 UV exposure. Once patterned, wells were profusely washed with PBS and then coated with fibronectin (10µg/ml). Six hundred cells were seeded per well for 1 h, then washed and the remaining cells were left to adhere for 12 hr.

### Western blotting

HUVECs were washed with cold PBS and scraped off in NETF buffer supplemented with protease and phosphatase inhibitors. Lysates were centrifuged and protein supernatant was quantified using the Lowry protein assay (Bio-Rad). Lysates were reduced in lamelli buffer and electrophoresis was performed using precast SDS-PAGE gels (Biorad) and transferred on to PVDF membranes (Biorad). Equal loading was checked using Ponceau red solution. Membranes were then blocked in 5% milk and probed with primary antibodies over-night (see below) and respective secondary antibodies (1:3000). The membranes were developed using ECL (Biorad) using the Chemidoc imaging system (Biorad). After initial immunodetection, membranes were re-probed with anti–GAPDH antibody. The protein bands were quantified using fiji and expressed as a relative value to either GAPDH or as the ratio of phosphorylated to total protein. The following antibodies were used: ARHGEF18 (1/500; GTX102223; Genetex), p38 (1:500; 9212, CST), Phospho-p38 (1:500; 9211, CST) ERK1/2 (1:500; 4695, CST), phospho-ERK1/2 (1:500; 9101, CST), Akt (1:500; 9272, CST), phospho-Akt (1:500; 9271, CST), Claudin5 (1:500; 34-1600, Invitrogen), RhoA (1:500; 2117, CST), Rac1 (1:500; 2465, CST), Vav (1:250; ab92890; abcam), GST (1:2000; PC53-100UG; Oncogene), paxillin (1:500; ab32084; abcam), phospho-paxillin (1:500; ab194738; abcam), GAPDH (1:1000; MAB374, MerckMillipore).

### In vitro Immunofluorescence staining

HUVECs were fixed with 4% PFA in PBS for 10 min at room temperature (RT). Blocking/ permeabilization was performed using blocking buffer consisting of 5% BSA (Sigma-Aldrich), 0.5% Triton X-100 (Sigma-Aldrich), 0.01% sodium deoxycholate (Sigma-Aldrich), and 0.02% sodium azide (Sigma-Aldrich) in PBS at pH 7.4 for 45 min at RT. Primary antibodies were incubated at the desired concentration in 1:1 Blocking buffer/PBS at RT for 2 hrs and secondary antibodies (Life Technologie, Alexa, 1/500) were incubated at the desired concentration in 1:1 blocking buffer/PBS for 1 hr at RT. DAPI (Sigma-Aldrich, 1/10000, 5 min) was used for nuclear labeling and phaloidin488 (life technologies, 1/500, 45min) was used to labelled polymerized actin. The following antibodies were used in vitro: VE-Cadherin (1/1000, RandD), ZO-1 (1/500, 61-7300, life technologies), claudin5 (MAB19903, Abnova, 1/500), paxillin (Abcam, 1/1000), FLAG (Merck, F1804), Donkey anti-Rabbit (life tech, 1/500), Donkey anti-Rat (life tech, 1/500), Donkey anti-Goat (life tech, 1/500), Donkey anti-Mouse (life tech, 1/500).

### In vivo Immunofluorescence staining

Retinas and Aortas were fixed with 4% PFA in PBS overnight at 4°C. Aortas were open “en face” before staining. Brains were sliced using a vibratome (100uM section). Blocking/ permeabilization was performed using blocking buffer consisting of 1% FBS, 3% BSA (Sigma-Aldrich), 0.5% Triton X-100 (Sigma-Aldrich), 0.01% sodium deoxycholate (Sigma-Aldrich), and 0.02% sodium azide (Sigma-Aldrich) in PBS at pH 7.4 for 1 hr at RT. Primary antibodies were used at the desired concentration in 1:1 Blocking buffer/PBS overnight at 4°C and secondary antibody were incubated at the desired concentration in 1:1 blocking buffer/PBS for 2 hrs at RT. DAPI (Sigma-Aldrich, 1/10000, 5 min) was used for nuclear labeling and Isolectin B4 488 (life technologies, 1/400, 1hr) was used to labelled endothelial cells. Retinas were mounted in Mowiol. The following antibodies were used in vivo: ARHGEF18 (HPA071867, 1/500), ERG (abcam ab92513, 1/1000), SM22 (abcam, 1/2000), TER119 (R&D MAB1125, 1/2000), VE-Cadherin (RandD AF1002, 1/500), ZO-1 (life tech 617300 1/1000), Donkey anti-Rabbit (life tech, 1/500), Donkey anti-Rat (life tech, 1/500), Donkey anti-Goat (life tech, 1/500).

### Impedance-based assay

Cell adhesion assay was performed using xCELLigence RTCA instrument (Roche). Briefly, HUVECs (P4-P6) expressing either shNT (non-target), sh18.1 or sh18.4 were plated in E-plate 96 well (5232368001; Agilent) at a density of 20,000cells/well. Cell adhesion was measured as electrical impedance between electrodes and represented as cell index value over a period of time.

### Wound healing assay

Confluent monolayer of HUVECs were scratched using a pipette tip. Cell debris were washed and the medium was replaced with fresh complete medium. Cell migration was video-recorded under optimal conditions (37°C and 5%CO2) using an inverted Ti Eclipse microscope equipped with a 10x/0.25 phase contrast objective and a DS-Qi2 CMOS camera (Nikon France, Champigny Sur Marne). Images were recorded every 5 minutes during 24 hrs. Wound area was manually measured at different time points using Fiji and rate of wound closure was calculated by area relative to time.

### Proliferation assay

Proliferation assay was performed using the Edu-click647 kit (BCK-EdU647; Base Click). HUVECs expressing the shRNAs were plated in μ-Slide VI 0.4 (80606; IBIDI) at a density of 30,000cells per channel in starving medium (EBM+0.5%serum+doxy+puro). After 16 hrs, cells were stimulated with complete medium (EBM+2%serum+doxy+puro) in the presence of EdU (5 uM) under static or flow for 8 hrs. Cells were fixed in 4%formaldehyde for 10 min, stained for EdU and nuclei were labeled with DAPI (62248; Thermofisher) according to the kit manual. DAPI+ and EdU+ cells were counted using Fiji.

### Microscope image acquisition

Images from fluorescently labeled HUVECs were acquired using a confocal A1 SIM microscope (Nikon) equipped with a Plan-Apochromat 20×/0.8 NA Ph2 objective or with a 60x/1.4 plan apochromat objective ; or using an Eclipse Ti2 inverted microscope (Nikon). Images were taken at room temperature using NIS Element software. Images of Retinas and Aortas were taken using a LFOV FLIM inverted confocal microscope (Nikon) equipped with a Plan-Apochromat 20×/0.8 NA Ph2 objective or with a Plan- Apochromat 60×/1.4 NA DIC objective. The microscope was equipped with a photon multiplier tube detector. Images were taken at room temperature using NIS element software (Nikon).

### Flow-induced orientation analysis

Cell shape was quantified using a dedicated FIJI macro. Basically, VE-Cadherin staining was detected in order to segment the border of each endothelial cell. Based on this segmentation a skeleton was built and aspect ratio (major axis lenght by minor axis lenght, AR) as well as the angle between the flow direction and the main axis was determined for all entire ECs visible on the image. Flow index corresponding to the ratio of AR and the angle was calculated for each EC. Flow index thus integrates both the elongation and the alignment of endothelia cells: The higher the flow index is, the better the cell response to the flow. Based on the VE-cadherin staining, gaps in the endothelial monolayer were quantified manually, only gaps with an area above 100um were taken in account.

### Cell junction activity analysis

ZO-1 intensity at the junctions was quantified using FIJI. Briefly, an enlarged mask from the VE-Cadherin segmentation previously described was generated and applied on the ZO-1 staining image, percentage of the ZO-1 staining in this area was extracted for each image. Actin cable formation was measured by the intensity of the phalloidin staining at the junctions using the same enlarged mask applied on the Actin staining image. Both analyses were made automatically by FIJI with identical parameter for mask enlargement and threshold values. Claudin5 localization at junction was quantified using a mix of Wildtype ECs and ECs expressing the shRNA. Using FIJI, intensity plots for both Claudin5 and VE-cadherin staining were obtained by drawing a line of approximately 15 um in between two cells.

Adherent junction morphology analysis was done in HUVECs or Aortas stained for VE-cadherin using the patch and classified Matlab code previously developed by Bentley and colleague^39^ adapted for 2D images. Two status were defined : activated vs stabilized, and divided into 3 level : from low to high. Activated junctions were defined as serrated or reticular and stabilized as straight think junctions. Each image taken was divided into small pieces of images allowing to visualized only a portion of the junction and these small images were presented blindly and randomly to the user who then classified the junction accordingly. This technic ensured an unbiased analysis of the junctions Irrespective of the treatment and entire shape of the cell.

### Basal adhesion analysis

For micropatterning images, paxillin channel was thresholded and the focal adhesion length was quantified for each focal adhesion detected. The focal adhesions were then classified by size within each cell. Quantification was done blindly.

For SS exposed ECs, focal adhesion type was assessed qualitatively by classifying blindly (by 2 operators) the overall aspect of Paxillin staining in the cell. Cells with long, numerous and parallel focal adhesions were classified 1 and cells with short, few and disorganized focal adhesions were classified 0.

### Statistical analysis

Statistical analysis was performed using GraphPad Prism software. For in vitro and in vivo experiments, when only 2 conditions were compared, a paired T-test or a Wilcoxon T-test were used depending on the distribution of the data. For multiple comparison an ordinary one-way ANOVA (data distribution was assumed to be normal) was used, followed by a Tukey test. Details of the statistical test used for each experiment can be found in the figure legends.

### Data Availability

The data that support the findings of this study are available from the corresponding author upon reasonable request.

## Acknowledgments

We thank Dr. Vincent Sauzeau and Dr. Thibaut Quillard for helpful comments on the manuscript. We are most grateful to the Genomics Core Facility GenoA, member of Biogenouest and France Genomique and to the Bioinformatics Core Facility BiRD, member of Biogenouest and Institut Français de Bioinformatique funded by the French National Agency (ANR) (ANR-11-INBS-0013) for the use of their resources and their technical support. We acknowledge the IBISA MicroPICell facility (Biogenouest), member of the national infrastructure France-Bioimaging supported by the ANR (ANR-10-INBS-04). This work was supported by ANR grants (Programme d’Investissements d’Avenir ANR-16-IDEX-0007 and ANR-21-CE14-0016 to A-C Vion) and the local fund Genavie (S. Batta). The authors declare no competing financial interests. Author contributions: S. Batta performed experiments, collected data, analyzed the results, and wrote the manuscript (original draft preparation). M. Rio, C. Lebot, C. Baron-Menguy, R. Moutaoukil performed experiments, collected data and analyzed results, R. Leruz performed experiments. G Loirand reviewed and edited the manuscript. A.-C. Vion conceived and designed the study, performed experiments, collected data, analyzed and interpreted the results, and wrote the manuscript. All authors reviewed the results and approved the final version of the manuscript.

