## Supplementary material for "ARHGEF18 is a flow-responding exchange factor controlling endothelial tight junctions and vascular leakage": Suplemental figures and tables

Supplemental Figure 1

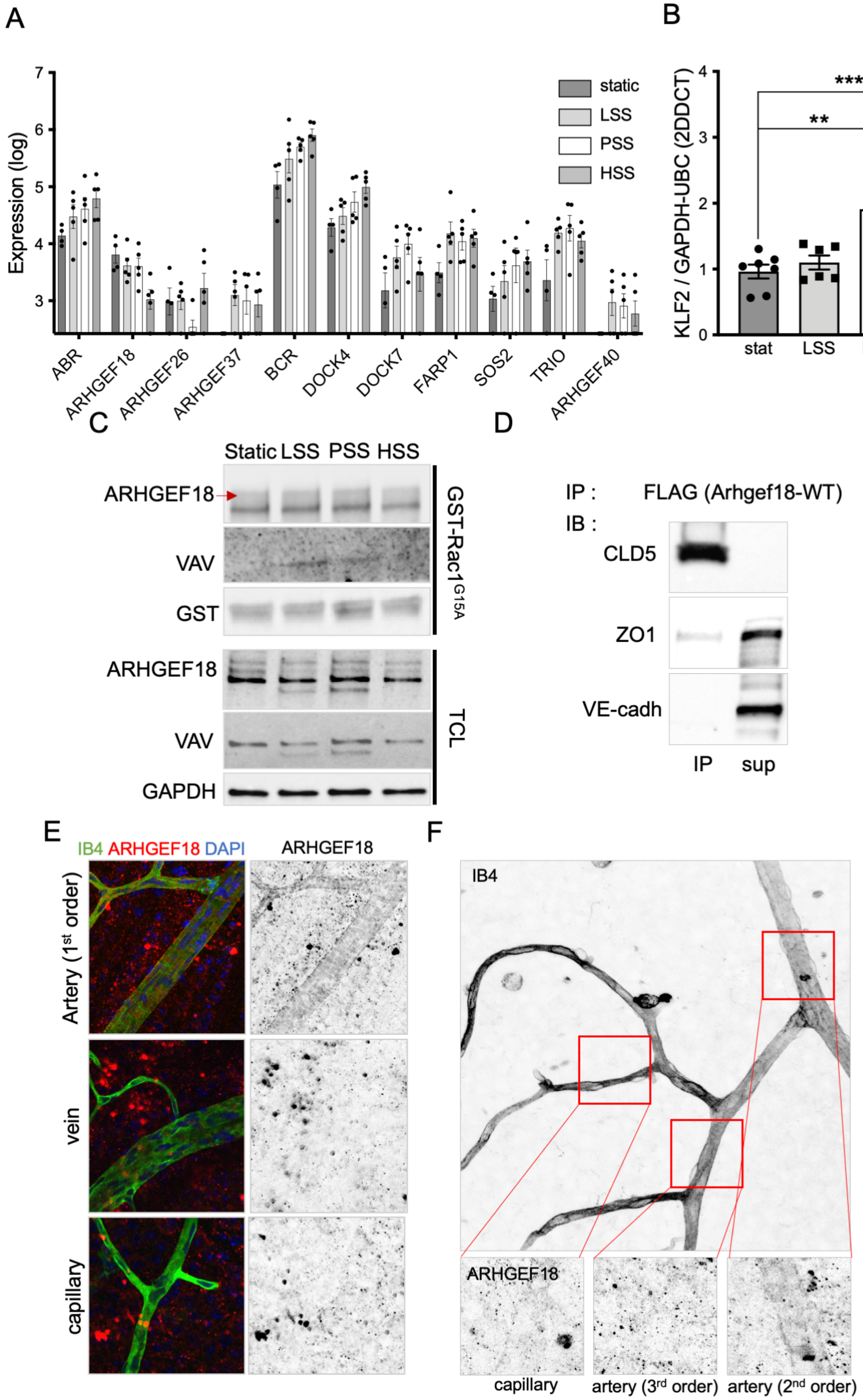

**Supplemental Figure 1: ARHGEF18 expression and activity is SS dependent. A.** extracted data form the 3'RNA sequencing analysis for the RhoGEFs with significant expression level changes. Static, N=4; SS N=5. LSS: low SS; PSS: physiological SS; HSS: high SS. **B.** Quantification of Klf2 expression level by qPCR. LSS: low SS; PSS: physiological SS; HSS: high SS. N>6. **p<0.01; ***p<0.001; One-way ANOVA**. C.** Representative blot of VAV binding on nucleotide-free Rac1 (GST-Rac1$^{G15A}$) by pull-down assay. 24h of SS, LSS: low SS; PSS: physiological SS; HSS: high SS. Representative of N=3. **D**. Representative blot from Co-immunoprecipitation of FLAG-ARHGEF18 with Claudin5, ZO-1 and VE-cadherin under static condition. Representative of N=2. **E.** Immunofluorescent staining of ARHGEF18 in 4-week-old mouse retina. Green: Isolectin B4; Representative of N=5. **F.** Second representative image from ARHGEF18 staining in the retina (4WO) showing ARHGEF18 staining in different area of the arterial tree.

### Supplemental Figure 2

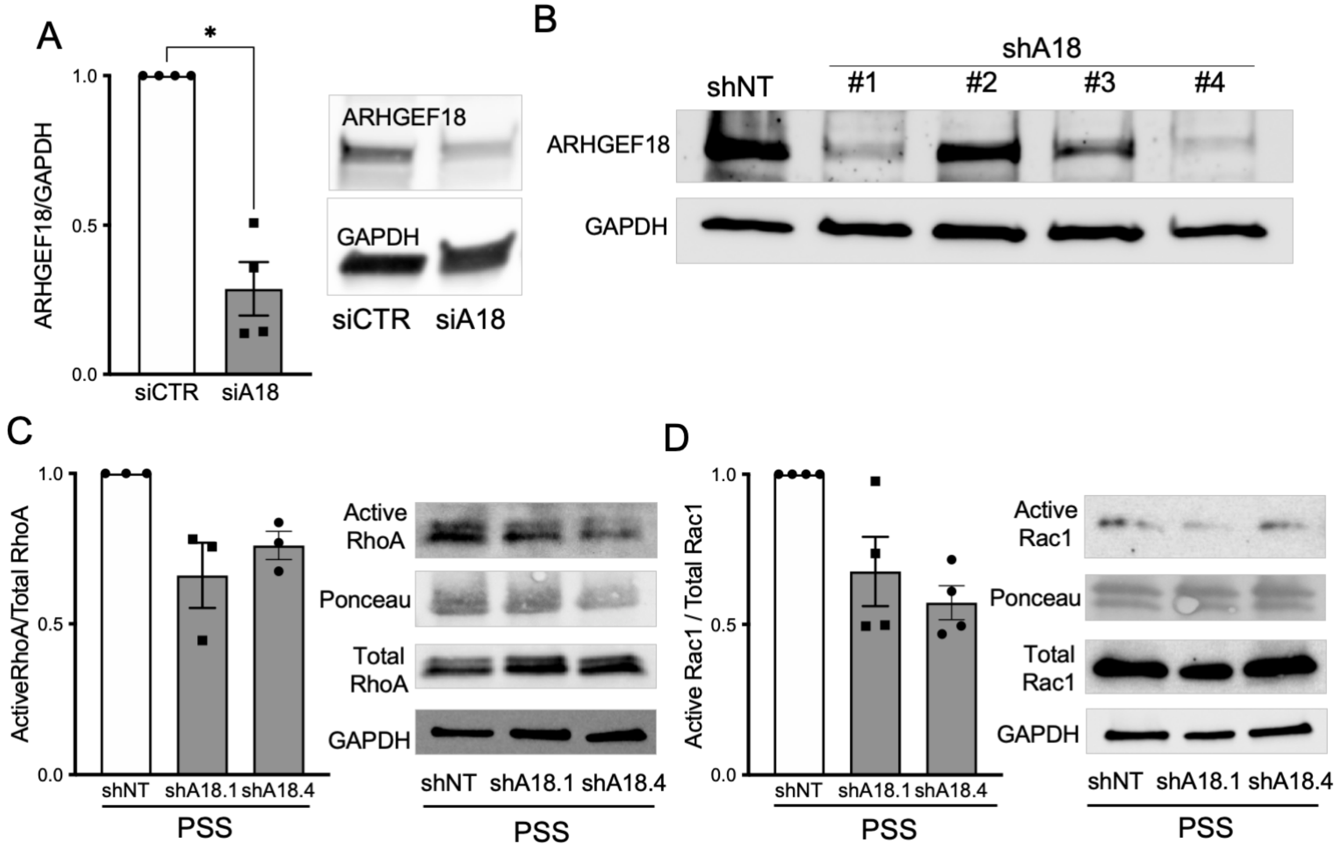

**Supplemental Figure 2: ARHGEF18 participates to endothelial cell adhesion and migration. A.** Silencing efficiency of the siRNA transfection assessed by Western Blot. N=4. Wilcoxon T-test. **B.** Representative Blot of the silencing efficiency by shRNA expression following infection, selection and induction (verified for each experiment done). **C.** Quantification of RHOA activity by pull-down assay using GST-RBD coated beads in HUVECs expressing a non-targeting shRNA (shNT) or a shRNA targeting Arhgef18 (shA18.1 or shA18.4) and exposed to physiological SS (PSS) for 24h. N=3**. D.** Quantification of RAC1 activity by pull-down assay using GST-PBD coated beads in HUVECs expressing a non-targeting shRNA (shNT) or a shRNA targeting Arhgef18 (shA18.1 or A18.4) and exposed to physiological SS (PSS). 24h of SS. N=4.

### Supplemental Figure 3

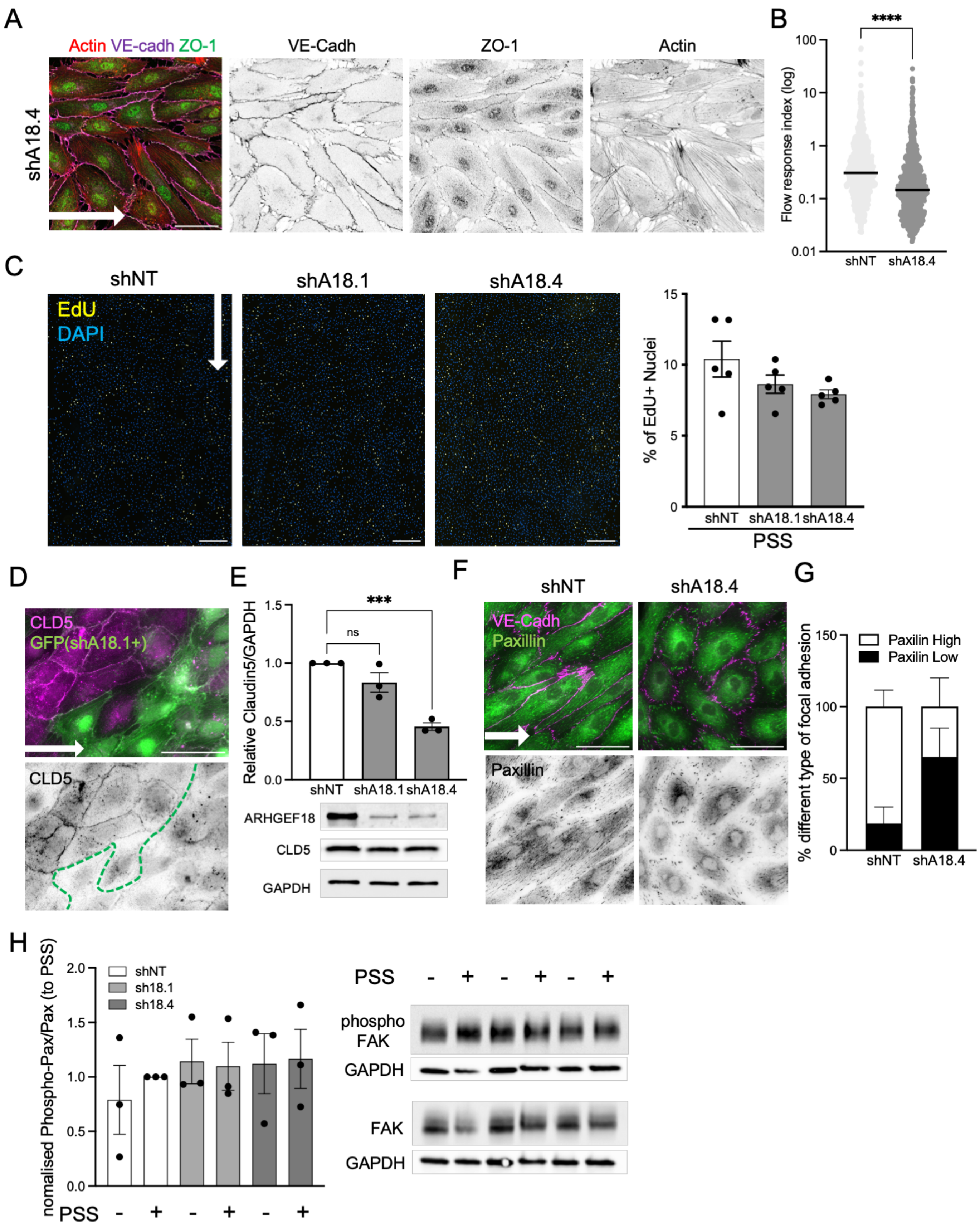

**Supplemental Figure 3: ARHGEF18 participates in ECs response to flow, tight junction formation and focal adhesion formation. A.** Representative images of HUVECs expressing a second shRNA targeting Arhgef18 (shA18.4), exposed to physiological SS and stained for ZO-1 (green), VE-cadherin (magenta) and actin (red). Flow direction is indicated by the arrow. Scale bar: 50μm. **B.** Quantification of cell orientation and elongation with the flow in HUVECs expressing a non-targeting shRNA (shNT) or a second shRNA targeting Arhgef18 (shA18.4), exposed to physiological SS for 24h. Each dot represents a cell. N=3 with in-between 200 and 350 cells analyzed by experiment. ***p<0.001; Mann-Whitney T-test. **C.** Representative images and Quantification of proliferation by Edu incorporation and staining in HUVECs expressing a non-targeting shRNA (shNT) or a shRNA targeting Arhgef18 (shA18.1 and shA18.4), exposed to physiological SS fro 8h. N=5. **p<0.01; 2-way ANOVA. Flow direction is represented by the arrow. Scale bar: 500μm **D.** Representative images of Claudin5 staining in a mix population of WT (GFP negative) and HUVECs expressing the Arhegf18 shRNA (shA18.1, GFP+), the green dotted line indicates the separation between WT and shA18 positive cells. Scale bar: 50μm. **E.** Quantification of Claudin5 protein level by Western blot in HUVECs expressing a non-targeting shRNA (shNT) or ARhgef18 shRNA (shA18.1 and shA18.4) exposed to physiological SS. N=3. ***p<0.001, One-way ANOVA. **F.** Representative images of Paxillin staining in HUVECs expressing a non-targeting shRNA (shNT) or a second Arhegf18 shRNA (shA18.4), exposed to physiological SS. Flow direction is indicated by the arrow. Scale bar: 50μm. **G.** Quantification of focal adhesion type by hand classification of paxillin staining within each cell based on the representative images for each category on the left (white box: paxillin high = numerous, elongated and aligned focal adhesions; black box: paxillin low = few, short and misaligned focal adhesions). N=2. **H.** Quantification of FAK phosphorylation level by Western blot in HUVECs expressing a non-targeting shRNA (shNT) or a shRNA targeting Arhgef18 (shA18.1 and shA18.4), under static condition or exposed to physiological SS. **I.** Classification and quantification of the junctional state of the adherent junction based on the morphological appearance with a VE-cadherin staining. N=4. One-way ANOVA. *p>0.05, **p<0.01.

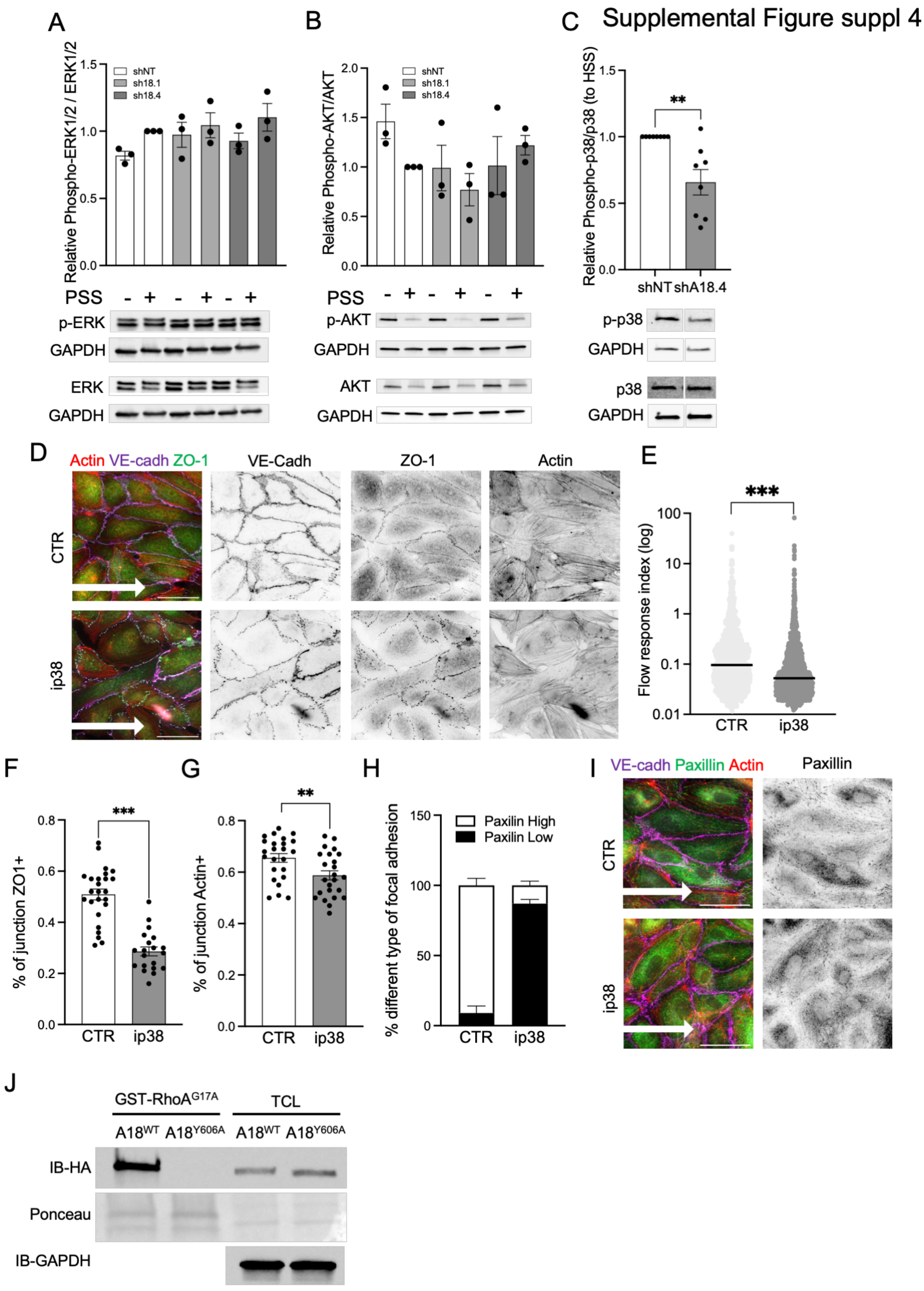
Supplemental Figure suppl 4
A
B
C
Relative Phospho-ERK1/2 / ERK1/2
shNT
sh18.1
sh18.4
PSS
p-ERK
GAPDH
ERK
GAPDH
Relative Phospho-AKT/AKT
p-AKT
AKT
Relative Phospho-p38/p38 (to HSS)
shNT
shA18.4
p-p38
p38
D
Actin VE-cadh ZO-1
VE-Cadh
ZO-1
Actin
CTR
ip38
E
Flow response index (log)
F
% of junction ZO1+
G
% of junction Actin+
H
% different type of focal adhesion
Paxilin High
Paxilin Low
I
VE-cadh Paxillin Actin
Paxillin
J
GST-RhoA^G17A
TCL
A18^WT
A18^Y606A
IB-HA
Ponceau
IB-GAPDH

**Supplemental Figure 4: ARHGEF18 effects under flow are dependent of p38 activity. A.** Quantification of ERK1/2 phosphorylation level by Western blot in HUVECs expressing a non-targeting shRNA (shNT) or Arhegf18 shRNA (shA18.1 and shA18.4), under static condition or exposed to physiological SS (PSS). N=3. **B.** Quantification of AKT phosphorylation level by Western blot in HUVECs expressing a non-targeting shRNA (shNT) or Arhegf18 shRNA (shA18.1 and shA18.4), under static condition or exposed to physiological SS (PSS). N=3. **C.** Quantification of p38 phosphorylation by Western blot in HUVECs expressing a non-targeting shRNA (shNT) or a second Arhgef18 shRNA (shA18.4), exposed to physiological SS. N=8. **p<0.01; paired T-test. Images originate from the same blot. **D.** Representative images of HUVECs treated with an inhibitor for p38 (ip38) or vehicle (CTR) exposed to physiological SS for 24 hrs and stained for ZO-1 (green), VE-cadherin (magenta) and actin (red). Flow direction is indicated by the arrow. Scale bar: 50μm. **E.** Quantification of cell orientation and elongation with the flow in HUVECs treated or not with an inhibitor for p38 (SB239063, 100nM, ip38), exposed to physiological SS for 24 hrs. Each dot represents a cell. N=4 with in-between 200 and 450 cells analyzed by experiment. ***p<0.001; Mann-Whitney T-test. **F.** Quantification of ZO-1 localized at junction in HUVECs treated or not with an inhibitor for p38 (ip38), exposed to physiological SS for 24h. N=2, 10 to 15 areas analyzed by experiment. ***p<0.001; Unpaired T-test. **G.** Quantification of Actin localized at junction in HUVECs treated nor not with an inhibitor for p38 (ip38), exposed to physiological SS for 24 hrs. N=2, 10 to 14 areas analyzed by experiment. **p<0.01; Unpaired T-test. **H.** Quantification of focal adhesion type by hand classification of paxillin staining within each cell based on the representative images for each category on the left (white box: paxillin high = numerous, elongated and aligned focal adhesions; black box: paxillin low = few, short and misaligned focal adhesions). N=2**. I.** Representative images of Paxillin staining in HUVECs treated or not with an inhibitor for p38 (ip38), exposed to physiological SS. Flow direction is indicated by the arrow. Scale bar: 50μm. **J.** Representative images of Western blot of ARHGEF18 binding on nucleotide-free RhoA (GST-RhoA$^{G17A}$) by pull-down assay with HUVECs exposed to physiological SS for 24 hrs, expressing a WT form (A18$^{WT}$) or a mutant form (A18$^{Y606A}$) of ARHGEF18.

### Supplemental Figure 5

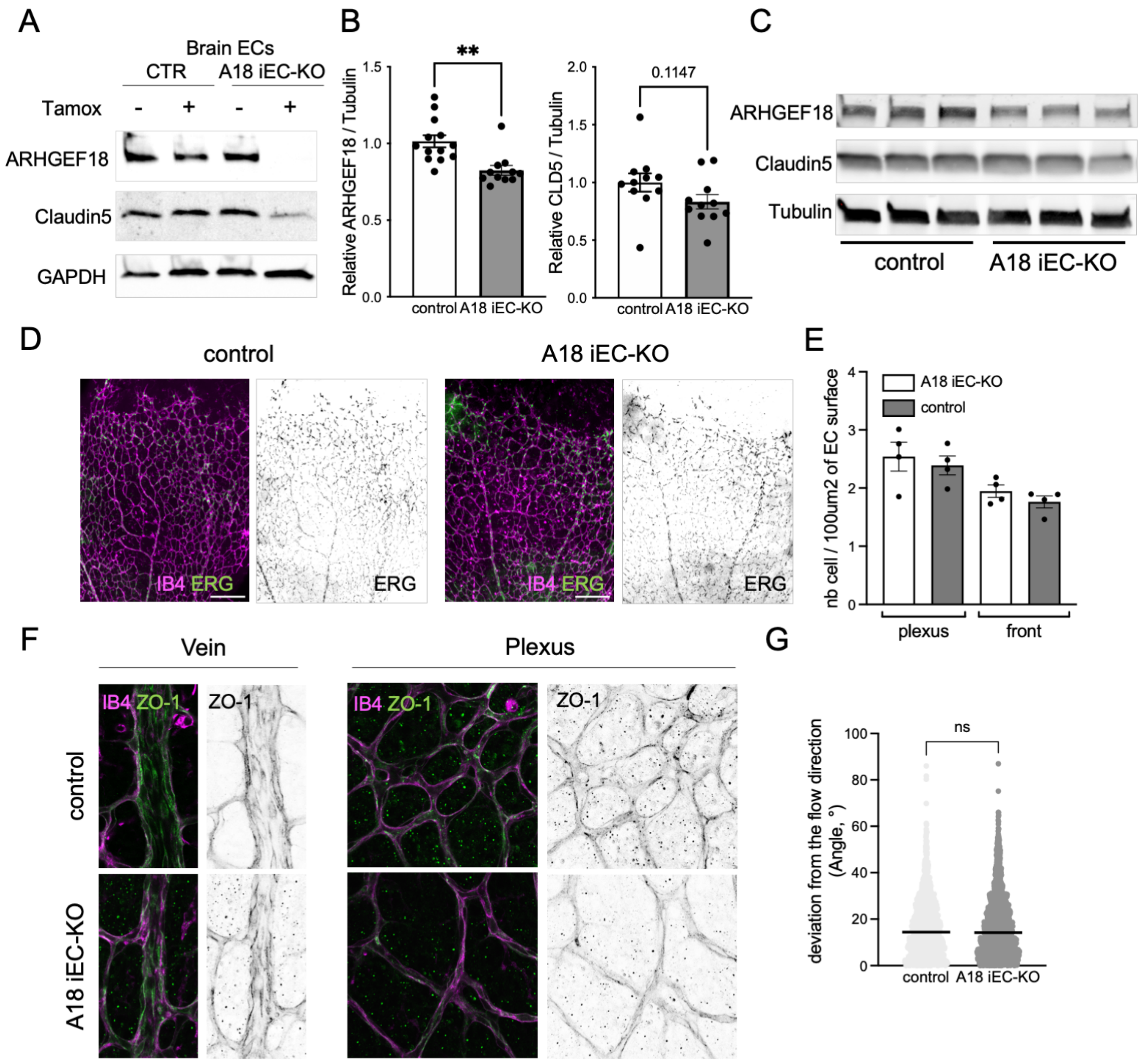

**Supplemental Figure 5: ARHGEF18 participates in vascular patterning and response to flow in vivo. A.** Representative blot of ARHGEF18 expression in brain endothelial cell extracted from *Arhgef18*$^{fl/fl}$*; CDH5-iCre*$^{ERT2+/wt}$ mice and littermate control (*Arhgef18*$^{fl/fl}$), treated or not with tamoxifen *in vitro*. **B.** quantification of ARHGEF18 and Claudin5 expression in lung of *Arhgef18*-iEC-KO mice and littermate control (P6), injected with tamoxifen (P1 to P3). N=13, Control; N=11, *Arhgef18*-iEC-KO. Mann-Withney T-test, **p<0.01. **C.** Representative blot of ARHGEF18 and Claudin5 expression in lung of *Arhgef18*-iEC-KO mice and littermate control. **D.** Representative image of ERG (green, black) and Isolectin (IB4, magenta) staining in the retinas of P6 pups. Scale bar: 200μm. **E.** Quantification of endothelial nuclei in *Arhgef18*-iEC-KO mice and littermate control (P6), in the plexus or invasive front areas. N=4. **F.** Representative images of ZO-1 (green, black) and Isolectin (IB4, magenta) staining in the vein or in the plexus of *Arhgef18*-iEC-KO mice and littermate control (P6) (observed in 3 littermate control and 4 *Arhgef18*-iEC-KO). **G.** Quantification of cell main axis angle compared with the flow direction in aortas. WT, N=6; *Arhgef18*-iEC-KO, N=7. Each dot represents a cell. in-between 200 and 300 cells analyzed by experiment.

#### Supplemental Figure 6

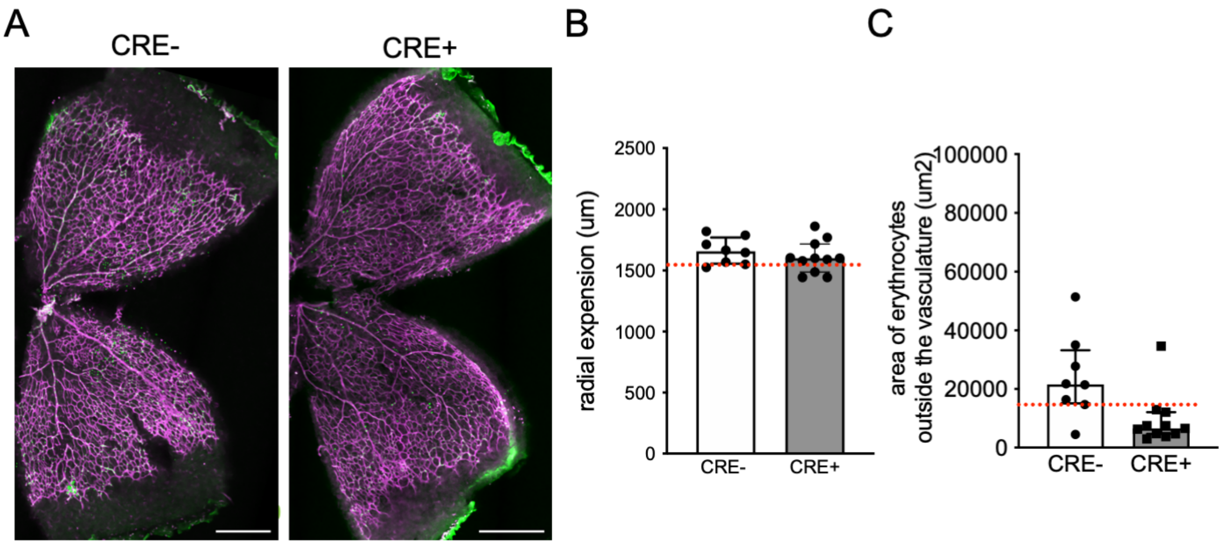

**Supplemental Figure 6**: **ARHGEF18 prevents retinas and brain hemorrhages. A.** Representative images of retinas from P6 CDH5(PAC)-CRE positive pups (CRE+) or littermate control (CRE-) stained with isolectin (IB4, magenta) and TER119 (green). Scale bar: 500μm. **B.** Quantification of the radial expansion of the vascular plexus in CDH5(PAC)-CRE positive pups (CRE+) littermate controls (CRE-, N=8; CRE+, N=11-; median ± IQ), red dotted line indicates the median value for *Arhgef18*$^{lox/lox}$ animals. **C.** Quantification of erythrocytes outside the vasculature in CDH5(PAC)-CRE positive pups (CRE+) littermate controls (CRE-, N=8; CRE+, N=11-; median ± IQ), red dotted line indicates the median value for *Arhgef18*$^{lox/lox}$ animals.

**Supplemental Tables**

Table 1: shRNA sequences used in this study

| Name | Sense (5’->3’) | Antisense (5’->3’) | Target |
|---|---|---|---|
| shNT (non-target) | CAAATCACAGAATCGTCGTAT | ATACGACGATTCTGTGATTTG | None |
| shArhgef18.1 | GGCCACAATGAAGCTGTTAGT | ACTAACAGCTTCATTGTGGCC | Exon |
| shArhgef18.2 | CAAAACTTGATCAAGAAAATT | AATTTTCTTGATCAAGTTTTG | Exon |
| shArhgef18.3 | CGGGCTACGACTGCACAAACA | TGTTTGTGCAGTCGTAGCCCG | Exon |
| shArhgef18.4 | AAGACAAGATGTCCTTTATGA | TCATAAAGGACATCTTGTCTT | Exon |
| shArhgef18.5 | CGATTTTATTTGTAAAGTTGA | TCAACTTTACAAATAAAATCG | 3’UTR |
| shArhgef18.6 | GAGCAAATGTTCCTATTTTCG | CGAAAATAGGAACATTTGCTC | 3’UTR |

Table 2: Primers and oligos used to amplify Arhgef18

| Oligo name | Oligo sequence (5’->3’) |
|---|---|
| Arhgef18_Fw | ACAGCTCTGCGATCGCCACCATGACGGTCTCTCAGAAAGGG |
| Arhgef18_Rev | GCTGTCTCGAGAATTAAGAAGAAGATGACGTCTTCTTTGC |
| ORF18-Oligo1 | ATGACGGTCTCTCAGAAAGGG |
| ORF18-Oligo2 | GTACGTCGGTCAGCAGGATAG |
| ORF18-Oligo3 | CTCACCTTCCGCAAGGAAGAC |
| ORF18-Oligo4 | GAAGTTGCGTTGCCGCTCCTG |
| ORF18-Oligo5 | CCCACCAGGACAGCTATGTG |
| ORF18-Oligo6 | GAAGAAGATGACGTCTTCTTTGCTGG |
